# Quantitative parameters of bacterial RNA polymerase open-complex formation, stabilization and disruption on a consensus promoter

**DOI:** 10.1101/2021.10.13.464252

**Authors:** Subhas C. Bera, Pim P. B. America, Santeri Maatsola, Mona Seifert, Eugeniu Ostrofet, Jelmer Cnossen, Monika Spermann, Flávia S. Papini, Martin Depken, Anssi M. Malinen, David Dulin

## Abstract

Transcription initiation is the first step in gene expression, and is therefore strongly regulated in all domains of life. The RNA polymerase (RNAP) first associates with the initiation factor *σ* to form a holoenzyme, which binds, bends and opens the promoter in a succession of reversible states. These states are critical for transcription regulation, but remain poorly understood. Here, we addressed the mechanism of open complex formation by monitoring its assembly/disassembly kinetics on individual consensus *lacUV5* promoters using high-throughput single-molecule magnetic tweezers. We probed the key protein–DNA interactions governing the open-complex formation and dissociation pathway by modulating the dynamics at different concentrations of monovalent salts and varying temperatures. Consistent with ensemble studies, we observed that RP_O_ is a stable, slowly reversible state that is preceded by a kinetically significant open intermediate (RP_I_), from which the holoenzyme dissociates. A strong anion concentration and type dependence indicates that the RP_O_ stabilization may involve sequence-independent interactions between the DNA and the holoenzyme, driven by a non-Coulombic effect consistent with the non-template DNA strand interacting with *σ* and the RNAP *β* subunit. The temperature dependence provides the energy scale of open-complex formation and further supports the existence of additional intermediates.

## Introduction

DNA-dependent RNA polymerase (RNAP) is the molecular machine responsible for the RNA production, i.e. the first step of gene expression, in cells (1,2). The structural core of RNAP is conserved in bacteria, eukaryotes and archaea (3–5). RNAP is a versatile target for antimicrobials having more than ten distinct small-molecule binding sites (6). Two antibiotics classes are currently in clinical use; rifamycins and fidaxomycin are used to treat tuberculosis and *Clostridium difficile*-associated diarrhea, respectively. The bacterial RNAP, typically consisting of five subunits (ααββ’ω), is structurally the simplest multi-subunits RNAP and has been widely employed as model system for molecular mechanisms of transcription and transcription regulation. The core RNAP cannot initiate promoter-specific transcription on its own. This deficit is compensated in bacteria by σ factors, which bind to the RNAP to form the transcription initiation competent RNAP holoenzyme complex (one σ per RNAP) (1,7).

Biochemical, structural and single-molecule studies have defined the basics of the transcription initiation process by bacterial RNAP–σ^70^ holoenzyme (from now on referred to as holo, and reviewed in (8,9)). This multistep mechanism begins with holo searching for promoters, embedded in the vast excess of non-promoter DNA. This search probably proceeds by a combination of modes, i.e. holo sliding along the DNA duplex (10–15), holo hopping from a DNA binding site to another nearby site (16) and holo diffusing through the bulk solution (17). The holo docks on the promoter via concerted interactions with specific promoter regions known as UP (from around −60 to −40 region of the promoter relative to transcription start site at +1 position), –35, spacer, and –10 elements (8,18–22). This initial unstable RNAP–promoter complex (RP_C_, where no DNA melting has occurred) isomerizes to more stable forms when the upstream and downstream regions of the promoter bend along the RNAP surface and into the DNA binding cleft, respectively (23–26). The formation of catalytically active holo–promoter open complex (RP_O_) is completed when the −11/+2 region of the promoter DNA duplex unwinds and the template DNA strand enters the active site cleft of the RNAP (27–30). The non-template (nt) DNA remains trapped outside the active site by the interactions between the –10 element and σ^70^ region 2, with −11A and −7T being flipped from the ntDNA base stack to deep pockets in σ^70^ (28,31). ntDNA binding is further stabilized by the discriminator (−6/+1) interactions with the σ^70^ region 2 and the core recognition element with the RNAP *β* subunit (27). Promoter sequence, transcription factors and small solutes modulate the stabilities and interconversion rates of the intermediates on the RP_O_ formation pathway and thus the level of gene expression (8).

Several studies have implied that not all formed RP_O_’s are structurally and functionally identical. Most studies report the existence of two (32–34) or more (35) open complex structures (RP_O_, intermediates), but also several closed complex intermediates have recently been identified (26). In most cases, less than half of apparent the RP_O_’s appear capable of productive promoter escape followed by full-length RNA synthesis (32,36–39). The majority of RP_O_ instead get trapped at the promoter and only synthesize short RNAs, and such moribund complexes (32) appear to play a role in transcription regulation (40). What these RP_O_ intermediates are, what interactions determine them, and what mechanisms allow their interconversion, remain poorly understood.

Single-molecule studies have revealed long-lived pausing and backtracking during transcription initiation (39,41). Instead of predefined RP_O_ populations producing different RNA types, a single RP_O_ could enter a pause/backtrack during initial RNA synthesis, before stochastically escaping the promoter. Controversy in data remains as a magnetic tweezers based single-molecule study reported only a single uniform RP_O_ (42), whereas a smFRET based study reported an additional low occupancy RP_O_ species with an unstable promoter conformation (43).

In the present study, we addressed the mechanism and heterogeneity of RP_O_ formation by monitoring the kinetics of RP_O_ formation and dissociation on individual DNA promoters using high-throughput magnetic tweezers. We probed the key protein–DNA interactions governing the RP_O_ pathway by modulating the dynamics with monovalent salts as well as temperature. We observed two different open conformations, i.e. one intermediate open complex RP_I_ and one final open state RP_O_. We show that the identity of monovalent cation mainly affects DNA twist, whereas the ranking of the anion in the Hofmeister series correlates with its effect on the transition from RP_C_ to RP_I_, indicating that this transition is driven by non-Coulombic interactions. Specifically, while physiological glutamate concentration favors rapid open complex formation, similar concentration of chloride promotes direct holo dissociation from the RP_I_ state to the degree that it does not populate the RP_O_ state anymore. From the strong salt dependence, we suggest that the stabilization of RP_O_ involves sequence-independent interactions between the DNA and the holo. One such candidate network of interaction takes place between the holo and the discriminator region of the promoter. Finally, the temperature dependence investigation revealed the energy landscape of open complex formation, the free energy difference between RP_C_ and dissociated holo, and supports the existence of several intermediate states during dissociation.

## MATERIALS AND METHODS

### High throughput magnetic tweezers

We used the high-throughput magnetic tweezers apparatus previously described in Ref. (44–46) to monitor dozens of individual DNA tethers in parallel. In short, it is a custom inverted microscope with a 50x oil immersion objective (CFI Plan Achro 50 XH, NA 0.9, Nikon, Germany), on top of which a flow chamber is mounted. The streptavidin coated magnetic beads (1 µm MyOne, ThermoFisher, Cat # 65001) are tethered to the bottom of the glass coverslip by a DNA construct that contains the *lacCONS+2* promoter for *Escherichia coli* RNA polymerase (see below *DNA construct fabrication*) (39) (**Figure 1**). A typical field of view is shown in **Supplementary Figure 1A** with hundreds of tethers and a few reference beads. An attractive force is applied to the magnetic beads to stretch the nucleic acid tether (**Figure 1A**) using a pair of vertically aligned permanent magnets (5 mm cubes, SuperMagnete, Switzerland). The magnets are separated by a 1 mm gap and are positioned above the objective as described in Ref. (44). The vertical position and rotation of the beads are controlled by the M-126-PD1 and C-150 motors (Physik Instrumente PI, GmbH & Co. KG, Karlsruhe, Germany), respectively. The field of view is illuminated through the magnets’ gap by a collimated LED-light source located above, and is imaged onto a large sensor CMOS camera (Dalsa Falcon2 FA-80-12M1H, Stemmer Imaging, Germany).

**Figure 1.**
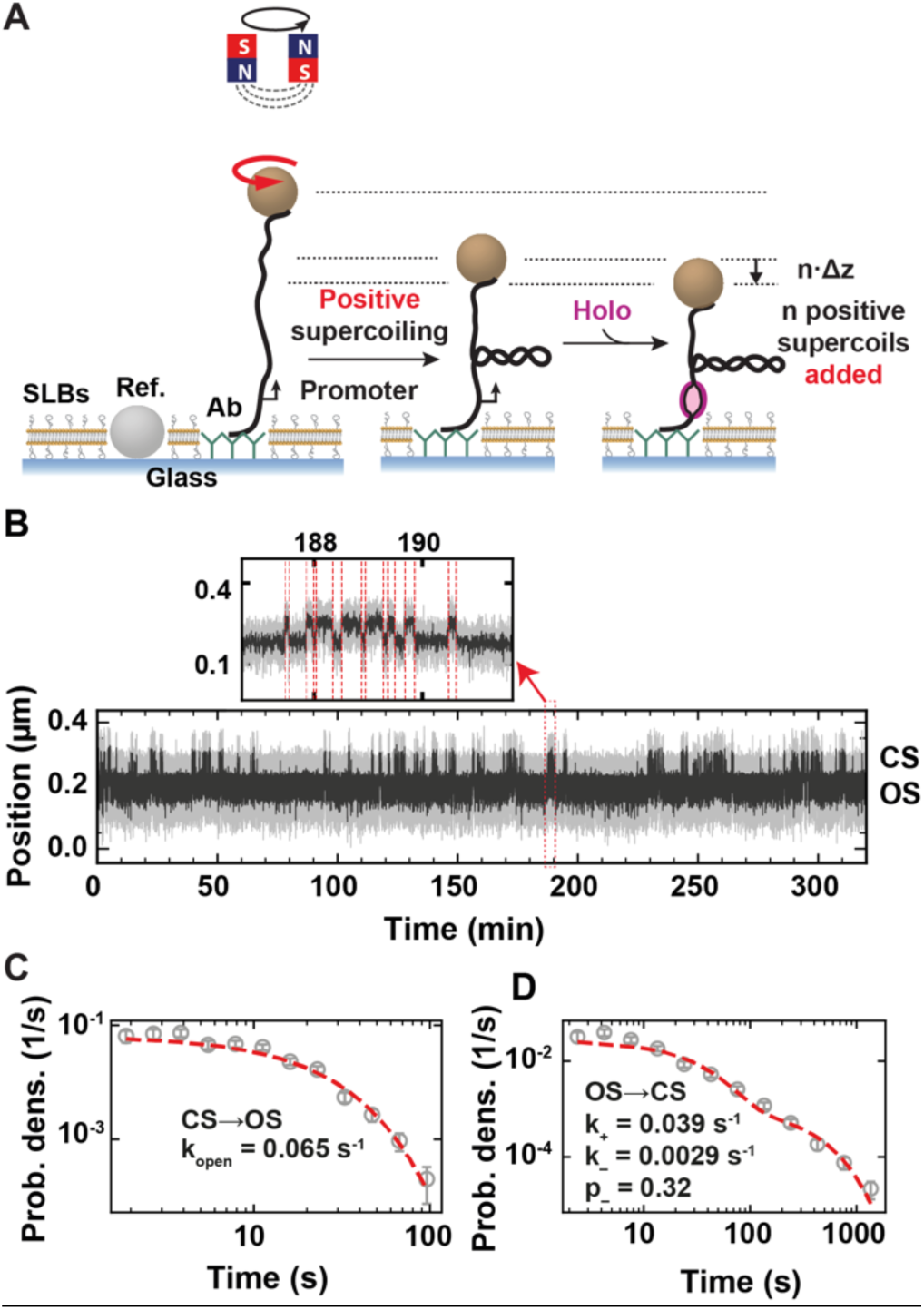
Measuring the bacterial holo open-complex dynamics using magnetic tweezers. **(A)** DNA plectonems formation upon positive supercoiling. Open complex formation by the holo leads to a decrease in the end-to-end extension of the DNA molecule by n.Δz, with n being the number of open base-pairs and Δz the distance by which the DNA extension decreases per full turn addition in a rotation extension experiment (**Supplementary Figure 1C**), i.e. Δz ~60 nm in reaction buffer. **(B)** Typical experimental trace showing dynamic transition between a promoter open and closed state (OS and CS, respectively) as described in (A). The experiment was performed with 10 nM holo, 150 mM KAc and at 34°C. A zoom in the trace shows fast open-complex dynamics. The red dashed lines indicate the transitions between the OS and CS captured by the change-point analysis (**Material and Methods**). **(C)** Probability density distribution of the CS dwell times for traces acquired as described in (B). The dashed red line is a maximum likelihood estimation (MLE) fit for a single exponential probability distribution function. The value of the mean exit rate *k*_*open*_ extracted from the MLE fit is indicated in the plot. **(D)** Probability density distribution of the OS dwell time distribution for traces acquired as described in (B). The dashed red line is a MLE fit for a double exponential probability distribution function. The values of the mean exit rate of the two exponentials, i.e. *k*_+_ and *k*_−_, and the probability *p*_−_ to exit the OS through the second exponential extracted from the MLE fit are indicated in the plot. Error bars in (C, D) are two standard deviations extracted from 1000 bootstraps.

### Preparation of vesicles

Small unilamellar vesicles (SUVs) were prepared by mixing DOPC (1,2-dioleoyl-sn-glycero-3-phosphocholine) and PEG-PE (1,2-dioleoyl-sn-glycero-3-phosphoethanolamine-N-[methoxy(polyethylene glycol)-550]) (850375C, 880530C, respectively, Avanti Polar Lipids, USA) dissolved in chloroform in 95:5 molar fraction. After mixing, the solution was dried under a stream of nitrogen, followed by drying in a vacuum desiccator for 1 h. The dry lipid film was then resuspended at 4 mg/mL in the storage buffer (10 mM HEPES, pH 7.8, 150 mM NaCl, 2 mM EDTA, 2 mM sodium azide) by vortexing. Then the suspension was incubated for 1 h at room temperature, with occasional vortexing and subsequently extruded 21 times through 0.1 µm polycarbonate membrane (Avanti, Cat # 610005) using a mini-extruder (Avanti Polar Lipids, USA, Cat # 610023). SUVs stock solution was stored at + 4 ºC for several weeks.

### Flow Cell assembly and lipid bilayer preparation

We used a lipid bilayer passivation strategy in the RP_O_ formation experiments, as our standard nitrocellulose passivation (44) failed to prevent non-specific sticking of the magnetic beads to the coverslip surface in the presence of holo. The glass coverslips used to assemble the flow cell (#1 thickness, 24×60 mm MenzelGlazer, Germany) were washed by sonication in a solution of Hellmanex III (Sigma-Aldrich, Germany) diluted in demineralized water (1% v/v). The coverslips were subsequently rinsed thoroughly under a stream of demineralized water, dried in an oven at 80°C, and stored in a 50 mL Falcon tube. A diluted stock of 1 µm polystyrene reference beads were prepared by diluting 1 µl stock (Sigma, Cat # LB11) 500-fold in mQ water, and rinsing them by repeating three times the following procedure: vortex, centrifugation at ~5000 rpm on a benchtop centrifuge for ~1 minute, removal of the supernatant and resuspension in the same volume of mQ water. The washed reference beads are finally resuspended in 50 µL mQ water and stored at 4°C for further use. ~4 µl of the reference bead solution, obtained by diluting the secondary stock 10-fold in absolute ethanol, was then spread with the side of a pipet tip on the top surface of the bottom coverslip of the flow cell, and subsequently heat up to ~130°C for ~2 minutes to melt the reference beads on the coverslip surface. Preceding the assembly of the flow cell, the side of the coverslips that forms the inner channel of the flow cell, were activated and made hydrophilic by thoroughly treating their surface with the electric discharge originating from a Corona SB (BlackHole Lab, Paris, France). The flow cell was then assembled by sandwiching a double layer of Parafilm (Sigma Aldrich, Germany, Cat # P7793) between two coverslips treated as described above. The flow cell was sealed by melting the Parafilm on a hot plate at ~100°C for 1 minute, while firmly pressing on top. The flow cell was subsequently mounted on the magnetic tweezers setup and rinsed with 1 ml 1x Phosphate buffered saline (PBS). 50 µl of full-length anti-digoxigenin (0.5 mg/mL in PBS, Sigma Aldrich, Germany, Cat # 11333089001) were added and incubated for 30 minutes. The excess was subsequently flushed away with 1 ml of 1x PBS buffer containing 700 mM NaCl followed by 10 minutes incubation, and a rinsing step with 1 ml PBS. The buffer in the flow cell was exchanged with 1 ml vesicle dilution buffer (10 mM HEPES at pH 7.4, 150 mM NaCl, 2 mM EDTA, 2 mM sodium azide and 2 mM CaCl_2_) to prepare the lipid bilayer assembly. The lipid bilayer was formed by flushing 1.2 ml of the SUV solution (see “*Preparation of vesicles”* below) at 50 µg/ml in the vesicle dilution buffer (Sigma Aldrich, Germany), slowly through the flow cell at 0.1 ml/min for a total duration of ~15 minutes. The flow channel was then washed with 1 ml PBS to remove any excess SUVs. To finalize the surface passivation, bovine serum albumin (BSA) (New England Biolabs, USA) at 1 mg/ml in PBS was incubated for 30 min, and subsequently flushed away excess BSA with 1 mL PBS.

In the meantime, 10 µl of MyOne streptavidin-coated superparamagnetic Dynabeads (Thermofisher, Germany, Cat # 65604D) were washed twice in PBS and diluted to 40 µL of PBS mixed with ~15 pM of DNA construct and 1 mg/ml BSA and incubated for a few minutes. The DNA tethered magnetic beads were then flushed into the flow cell and incubated for ~5 minutes to ensure attachment of the digoxygenin-labelled DNA handle to the anti-digoxigenin adsorbed on the flow cell surface. Finally, the excess of magnetic beads was removed by flushing copious amounts of PBS with occasional gentle tapping on the exit tube connecting the flow cell to the withdrawing pump.

### DNA construct fabrication

The long linear DNA construct (20666bp) was obtained by plasmid digestion and ligation with functionalized digoxigenin-or biotin-handles (850bp), obtained from PCR on λ DNA (44). The fabrication of the 1.4kb DNA construct was done as described in Ref. (47). Briefly, the desired DNA fragments were amplified by PCR from a synthetic plasmid containing the *E. coli* RNAP *LacCONS* promoter sequence and selectively cleaved by nicking enzymes at multiple sites to obtain ssDNA of different lengths and partial complementarity. These fragments were annealed to form a double strand with functionalized biotin and digoxygenin handles. The resulting nicks were ligated to obtain a torsionally constrained molecule.

### DNA tether selection

We first evaluated the extension and coilability of the DNA molecule by stretching the DNA from zero to 4 pN and rotating the magnetic bead first in negative and then in positive directions (± 15-100 turns depending on the length of tether), while applying 4 pN force. For a coilable and single-tether DNA molecule, the extension should remain constant in the negative turn region but decrease in the positive turn region (48).

### DNA Rotation experiment in different monovalent salts

To determine the effect of monovalent salt on DNA twist, we used a coilable ~20.6 kbp DNA molecule, which fabrication was described elsewhere (49,50). After selection of coilable DNA molecules, the flow cell was flushed with a reference buffer (10 mM Tris-HCl, 150 mM NaCl, 2 mM EDTA) with occasional mild tapping (without mounting the magnets) to relax the DNA from any unwanted twist. The rotation experiments were performed by slowly rotating the magnets (2 turns/sec) from −70 to 70 turns at ~0.3 pN force, the zero twist in the tethers being defined by the maximum extension in the reference buffer. The buffer (10 mM Tris_HCl, 2 mM EDTA) containing the monovalent salt of interest was flushed in the flow cell while applying a high force to the DNA (8 pN) and at zero turn. After 3 minutes of incubation, the same rotation experiment was performed in the experiment buffer containing the monovalent salt. Finally, the flow cell was flushed with the reference buffer while clamping the DNA with high force (8 pN) and after 3 minutes of incubation, another measurement was performed in the reference buffer. The first and the last reference measurement were performed to check whether the DNA supercoiled state had been restored in reference buffer to eliminate tethers in which a change in twist may have occurred from the magnetic bead sticking to the surface. When changing salt condition, the flow cell was flushed in with the reference buffer without the magnet to allow the DNA to relax and the same procedure was repeated. These experiments were performed at 25 °C.

### Magnetic tweezers holo open complex dynamics experiments

To monitor the open complex dynamics, we tethered magnetic beads with the ~1.4 kbp coilable DNA molecules. The flow cell was rinsed with reaction buffer (40 mM HEPES, 10 mM MgAc_2_, 1 mM DTT, 1 mM cysteamine hydrochloride, 5% glycerol, pH 7.8 and the monovalent salt of interest at the indicated concentration) before the addition of the enzymes. The buffer was always pre-heated to the experiment temperature, i.e. 34°C if not indicated otherwise, before injecting into the flow cell. Preceding the measurement, we evaluated that the DNA tethers were relaxed. To this end, following data acquisition, DNA force-extension (varying the applied force from zero to high) and rotation-extension (twisting the tethers from −5 to 5 turn at 0.3 pN) tests were performed before holo addition in the flow chamber at +3 turns and 8 pN force (such force was used to prevent tethers twist change and magnetic bead sticking while flushing). The force was subsequently reduced to 0.3 pN and data acquisition was continued until the end of the experiment.

### Dwell times detection

To identify the points at which the DNA transitions between states (opened and closed state), the so-called change points, we used an offline change point detection. With this analysis method, change points in the recorded data can be identified through finding the abrupt changes in the average magnetic bead position, even though neither the location nor the number of break points were known. For our purpose, we work with the change point detection algorithm as implemented in the Python package Ruptures (51). Change point detection is defined by a search method, a cost function and a constraint. The search method defines the algorithm used to analyze the time series. Here, as the true number of change points is unknown, we use the Bottom-Up algorithm. A dataset of length n is separated into n/2 segments and the pair segments with the lowest cost are merged until crossing a user defined penalty (see below). Bottom-Up has been shown to outperform the Binary Segmentation algorithm (52). As a cost function, we used a least absolute deviation to detect the changes in the median position of the magnetic bead. With an unknown number of change points, a constraint (penalty) is needed to balance out the goodness of fit parameter. The penalty was determined by manual inspection of change point detection quality for a particular data set. We use a penalty between 0.2 and 5 depending on the durations of the states. We embedded the Ruptures package into a GUI that is provided in our lab GitLab account (https://gitlab.com/DulinlabVU/change_point_analysis).

### Maximum likelihood fitting

The procedure is described in detail in Ref. (39). Briefly, the dwell times *τ* are described according to a probability-distribution function consisting either one or two exponentials,

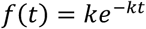

and

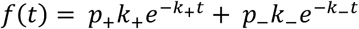

respectively. Here *k, k*_+_, *k*_−_ are characteristic rates, *p*_+_ and *p*_−_ are probabilities normalized such that *p*_+_ + *p*_−_ = 1. The number of exponential fit to the data was determined using the Bayes Schwarz Information Criterion (BIC) (53). We calculate the maximum likelihood estimate of the parameters (MLE) (54) by minimizing the negative of the likelihood function

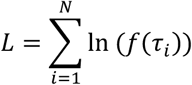

over the parameter set using minimization function with ‘L-BFGS-B’ algorithm in SciPy (SciPy.org). Here the *τ*_*i*_ are the experimentally measured dwell times and *N* is the number of collected dwell times *τ*_*i*_. The one standard deviation statistical error were extracted for each fitting parameter from 1000 bootstraps (55).

### Kinetic description of the open state formation

We consider the formation of the first state of the open complex, i.e. RP_I_, to be governed by the reaction scheme

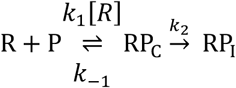

Previous studies have shown that holo binding equilibrates rapidly, i.e. with an association equilibrium constant *K*_1_ = *k*_1_/*k*_−1_, in comparison to the slow isomerization from RP_C_ to RP_I_ with the reaction rate constant *k*_2_ (8) (**Figure 1E**). We therefore describe the observed rate of OS formation *k*_open_ as (56)

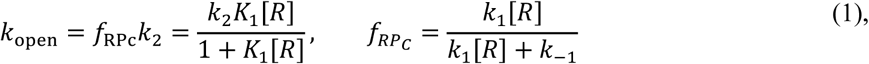

which coincides with the ensemble description of Ruff and co-workers (8). In the above, *f*_RPc_ is the fraction of promoter at equilibrium in the RP_C_ state.

### Kinetic description of the OS to CS transition

The dissociation of the holo from the promoter is described by the kinetics of the transition from OS to CS. After evaluating several kinetic models to describe the OS dwell time distribution (**Table 1**), we concluded that the best model corresponds to Model 3, Assumption 3 (**Table 1**):

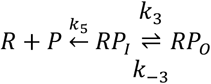

**Table 1:**
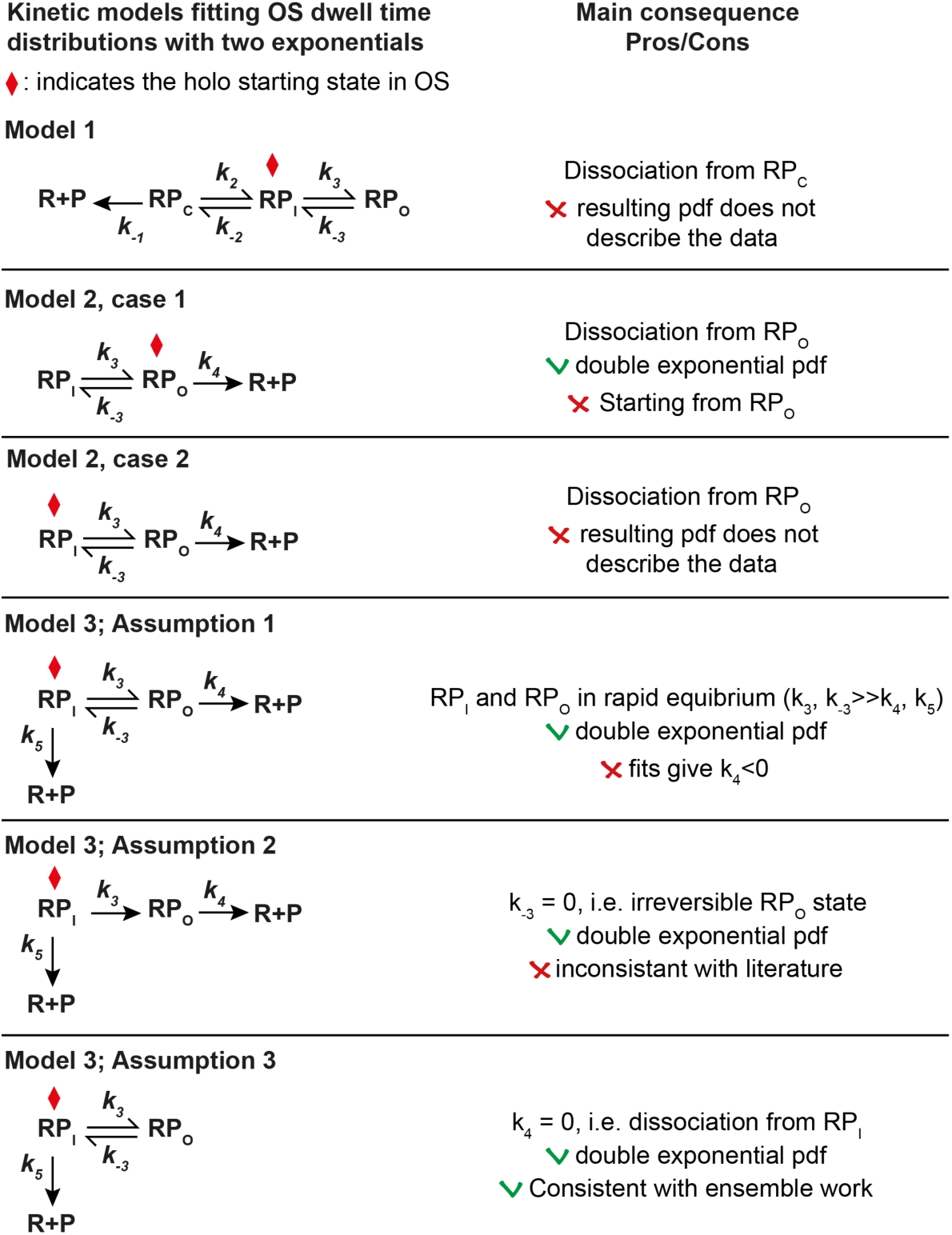
Comparison of models describing a double exponential distribution of the OS dwell times. pdf: probability distribution function. The left column presents three models. Model 1 describes a reversed kinetic pathway to dissociation from RP_C_. Model 2 hypothesizes a dissociation from RP_O_, with two specific cases, i.e. the holo starts in OS from either RP_O_ or RP_I_, and eventually dissociates from RP_O_. Model 3 includes dissociation from RP_I_ and is solved in the context of three assumptions for which we have an analytical solution. The right column indicates the key parameter of the model, and the reason to either discard or keep a specific model.

Under this assumption, we obtained a complete set of conversion relations for the fit parameters to the kinetic rates (**Supplemental information**).

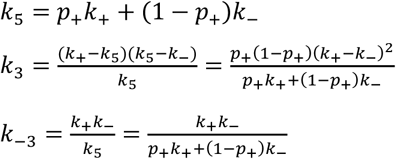

Note that since 0 ≤ *p*_+_ ≤ 1, all the microscopic rates are positive, as should be.

### Error propagation from the double exponential fits to the kinetic rates

We propagated the errors of the MLE fitting parameters (Δ*k*_est,*i*_) to the rate constant of the kinetic model (**Figure 1E**) as

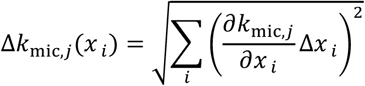

where *k*_mic,*j*_ is either of the derived microscopic rates *k*_4_, *k*_−3_, or *k*_3_, and *x* _*i*_is either the estimated rates or probabilities *k*_+_, *k*_−_ or *p*_−_. We use one standard deviation statistical error extracted from the bootstrap procedure.

### Theoretical description of the temperature dependence of the kinetic rate constants

For a two-state transition, the temperature dependence of the forward reaction is empirically described by the Arrhenius equation

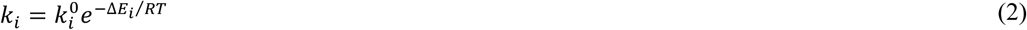

where 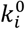 is the exponential prefactor, *E*_*i*_ is the activation energy for the reaction rate constant *k*_*i*_, *R* is the gas constant, and *T* is the temperature. **Equation 2** describes well *k*_4_, *k*_3_ and *k*_−3_ (**Figure 6CDE**).

We used here the model represented in the above section to describe the opening state formation rate, i.e. *k*_open_. From this model, we have at equilibrium (57)

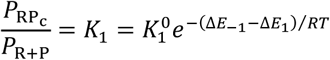

Where 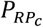 and *P*_*R*+*P*_ are the probabilities to be in the RP_C_ or the R+P (dissociated) states, respectively, 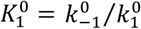 is the dissociation constant at 25°C, Δ*E*_−1_ and Δ*E*_1_ are the activation energies of the transition towards either dissociation or association, respectively. Using **Equation 1** with the rate constants described by Arrhenius (**Equation 2**), we obtain

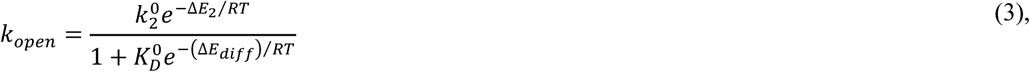

where Δ*E*_*diff*_ = Δ*E*_−1_ − Δ*E*_1_, Δ*E*_2_ is the activation energy of the transition from RP_C_ to RP_I_, 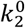 is *k*_2_ at 25°C and 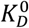 the dissociation constant at 25°C. To have the fit converging, we fit out the natural logarithm of the rates, and then converted them back. We provided initial parameters values for the fit by fitting **Equation 2** to the two regimes observed in **Figure 6B**, from which we extracted *E*_2_ = (20 ± 2) kcal · mol^−1^ and *E*_−1_ = (−63 ± 3) kcal · mol^−1^ for temperatures below and above 37°C, respectively (blue and red dashed lines, respectively **Figure 6B**).

## RESULTS

### High-throughput magnetic tweezers assay to study open complex dynamics for the bacterial RNA polymerase

We used a high-throughput magnetic tweezers assay to monitor the open complex dynamics for experiments up to several hours (**Figure 1A, Supplementary Figure 1A**). The magnetic bead was tethered to the top glass surface of the bottom coverslip of a flow cell by a ~1.4 kilo-basepair (kbp) torsionally constrained DNA molecule, i.e. without nicks and with multiple attachment points between the handles and either the magnetic bead or the glass surface (**Figure 1A**) (**Material and Methods**). The flow cell surface was passivated using a lipid bilayer strategy, which significantly reduces the non-specific adhesion of magnetic beads and proteins to the flow cell surface (58–61). We provide a detailed protocol in the **Materials and Methods** section to establish such passivation strategy. The DNA sequence encodes the *lacCONS+2* promoter for *Escherichia coli* holo (**Supplementary Figure 1B**), which is a consensus versions of the *lacUV5* promoter (39), and has been extensively studied in ensemble (8) and single molecule experiments investigating bacterial transcription initiation (62).

The rotation of the magnets above the flow cell induces rotation of the magnetic bead, and concomitantly adds twist in the DNA tether. As the number of turns in the DNA molecule increases, torque also increases, up to the buckling transition at which the DNA starts forming plectonemes (48,63,64). Subsequent addition of turns to the DNA molecule is then converted into writhe, while keeping the torque constant, leading to a decrease of the DNA molecule end-to-end extension (48). In a rotation-extension experiment at ~0.3 pN, the DNA extension is maximal at zero turn and decreases symmetrically when adding either positive or negative turns (**Supplementary Figure 1C**) (48). The linking number is the conserved sum of writhe and twist in a torsionally constrained nucleic acid molecule (65). The opening of the promoter (i.e. DNA unwinding or bubble formation) by the holo reduces the twist of the molecule, which is compensated by increase in writhe for a positively supercoiled DNA molecule. This leads to a decrease of the end-to-end extension of the DNA molecule by *n* · Δ*z*, with *n* being the number of open base pairs, and 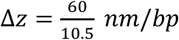 (for a DNA helical pitch of ~10.5 bp/turn) the rate at which the DNA molecule extension decreases per added twist (**Figure 1A, Supplementary Figure 1C**), as described by Strick and colleagues (42).

For a positively supercoiled DNA molecule and in the presence of holoenzyme, we clearly distinguished two main magnetic bead vertical positions, i.e. one indicating a DNA molecule without open complex that we coined the closed state (CS) and another one reporting a shorter DNA molecule end-to-end extension that signals a formed open complex and we coined the open state (OS) (**Figure 1B**) (42). The CS dwell time is the total time required for the holo to find, bind and open the promoter, while the OS dwell time is the total time the promoter is open until closing. We investigated only the positive supercoils regime, as the open complex with *lacCONS* promoter is so stable in the negative supercoil regime that it does not close back once open, i.e. showing no dynamics (42). As torque influences the open complex dynamics (42), we have only selected DNA molecules that were in a torsional state in the linear regime following the buckling transition in a rotation-extension experiment, where the torque is constant (**Supplementary Figure 1C-F**). A visual inspection of a magnetic tweezers trace shows short-lived CS interrupted by either short or long-lived OS (**Figure 1B**). To quantify our observation, we use a Python-written custom graphic user interface (GUI), based on the change-point algorithm Ruptures (51) (**Material and Methods**, provided in https:/gitlab.com/DulinlabVU/change_point_analysis), to automatically detect the transitions between OS and CS states in the magnetic tweezers traces. The extracted CS and OS dwell times were subsequently assembled in distributions, which were fitted using a maximum likelihood estimation (MLE) procedure (54). The CS dwell time distributions were best fitted by a single exponential probability distribution function with a promoter opening rate *k*_*open*_ (**Figure 1C, Materials and Methods**). In the presence of 150 mM potassium acetate (KAc), the OS dwell time distribution was best fitted by a double exponential probability distribution function with the fitting parameters *k*_+_, *k*_−_ and *p*_−_, i.e. the characteristic rates of the first and second exponential, respectively, and the probability of the second exponential (**Figure 1D**) (**Materials and Methods**) (39). This double exponential distribution was not reported in previous magnetic tweezers study of the OS dynamics (42).

### A rapidly equilibrating binding followed by promoter opening describes open state formation

We first investigated the formation of open complex in 150 mM KAc at different holo concentrations. A direct observation of the traces shows a shorter CS dwell time when increasing holo concentration from 0.2 to 2 nM (**Figure 1B, Figure 2A**). Extracting *k*_*open*_ from the CS dwell time distributions (**Supplementary Figure 2**), we found *k*_*open*_ to increase with holo concentration (**Figure 2B**) and be well described by a model where holo association to/dissociation from the promoter equilibrates quickly in comparison to the isomerization from RP_C_ to RP_I_ (**Equation 1, Figure 2C**), supporting previous magnetic tweezers observation by Strick and colleagues (42). From fitting **Equation 1**, we extracted the equilibrium binding constant *K*_1_ = (60 ± 5) µ*M*^−1^ and the rate constant *k*_2_ = (0.17 ± 0.01)*s*^−1^ (**Table S3**). The latter is only approximative, as the assay temporal resolution could not saturate the holo binding kinetics. The CS kinetics are consistent with a complete dissociation of the holo from the promoter following the transition from OS to CS, and the binding of a different holo preceding the next OS formation. This is further supported by the following three experiments. We first incubated the flow chamber with 0.5 nM holo in a reaction buffer to initiate and record CS–OS dynamics. The flow chamber was flushed about ~1800 s after the start of the recording either with reaction buffer (**Supplementary Figure 3A**), reaction buffer containing ~10 nM competing *lacCONS+2* DNA promoter fragment (**Supplementary Figure 3B**) or reaction buffer containing 100 µg/ml heparin (**Supplementary Figure 3C**). All three experiments showed that after the flushing step the CS never converted back to the OS (**Supplementary Figure 3**). This finding confirms that transition from OS to CS leads to the full release of the holo from the promoter.

**Figure 2.**
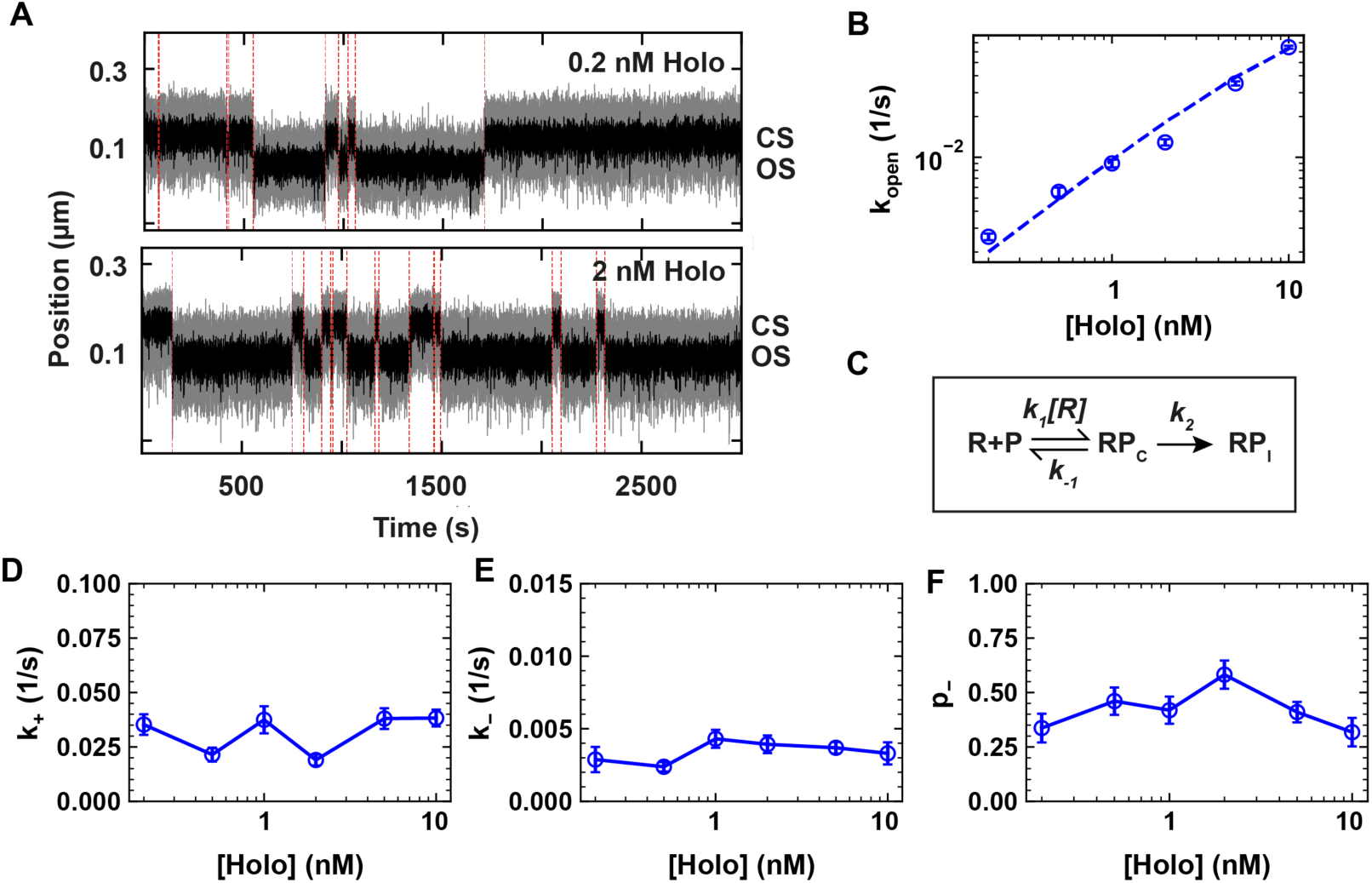
The CS dynamics is consistent with holo dissociation at the transition from OS to CS. **(A)** Traces showing the RP_O_ dynamics at 0.2 and 2 nM RNAP concentrations in 150 mM KAc. The dashed red lines indicate the transition between CS and OS. **(B)** Kinetic model of the OS formation. RP_C_ and RP_I_ respectively indicates the holo-promoter closed complex and the holo-promoter intermediate open complex. *k*_1_, *k*_−1_, and *k*_2_ are the reaction rate constants for holo association, dissociation, and open-intermediate. **(C)** CS exit rates *k*_*open*_ as a function of holo concentration. The dashed line is a linear fit of the model described in (B) (**Equation 1, Materials and Methods**). **(D, E, F)** *k*_+_, *k*_−_ and *p*_−_ as a function of holo concentration using the reaction buffer containing 150 mM KAc. Error bars are one standard deviation extracted from 1000 bootstraps in (C, D, E, F).

**Figure 3.**
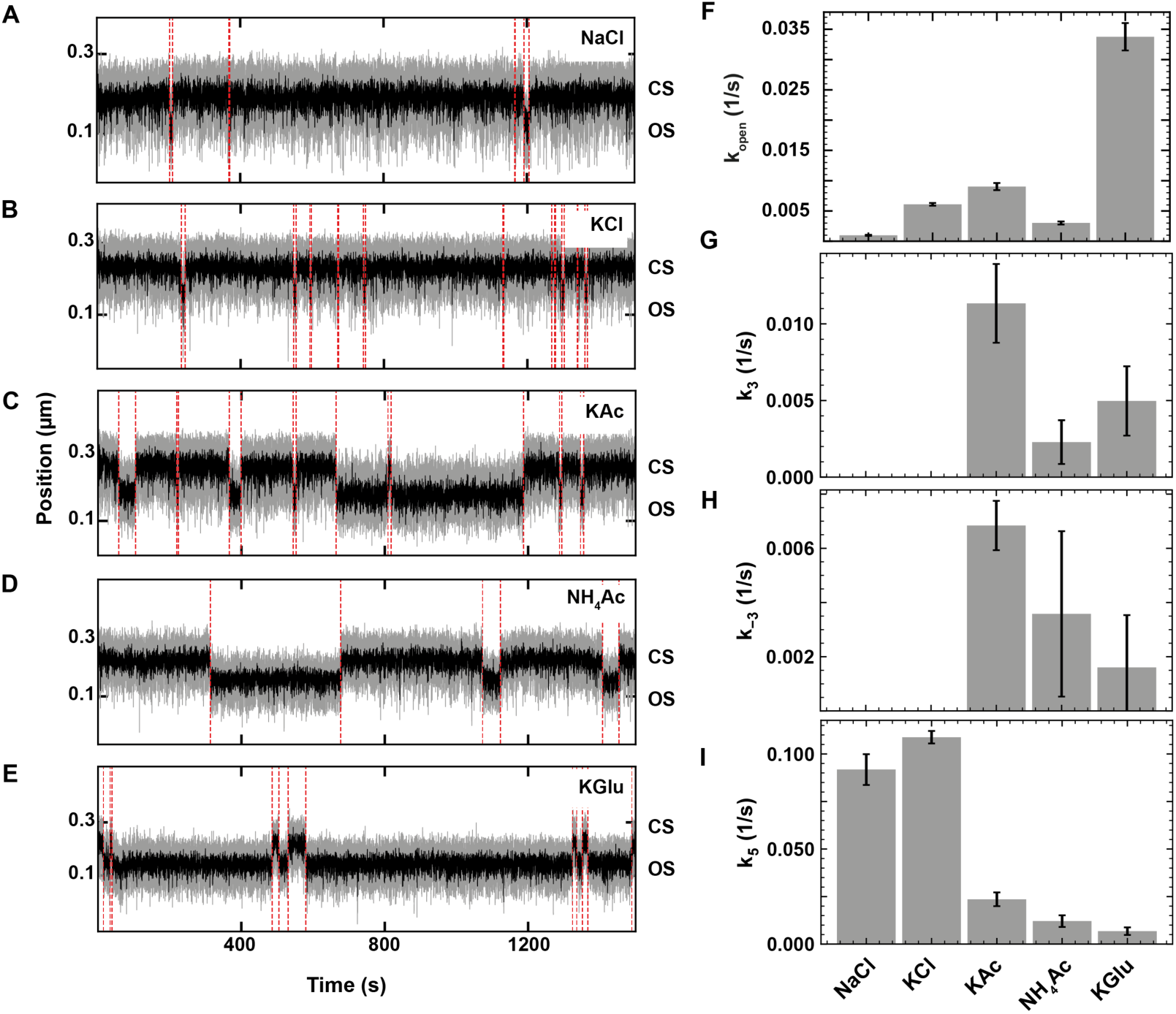
Monovalent salts affect bacterial holo open complex dynamics. **(A, B, C, D, E)** Holo open complex dynamics was observed at 34°C in the presence of 150 mM of the indicated monovalent salt. 10 nM holo was used in KCl and NaCl (A, B), and 1 nM holo was used in KAc, NH_4_Ac and KGlu (C, D, E). The red dashed lines indicate the transitions between OS and CS captured by the change-point analysis. **(F, G, H, I)** Dependence of *k*_*open*_, *k*_3_, *k*_−3_ and *k*_5_ on the monovalent salt type (Model 2, Assumption 3 in **Table 1**). The second exponential was absent from the dwell time distributions when using either NaCl or KCl, and we therefore extracted only *k*_5_ for these conditions. Error bars in (F, G, H, I) are the propagated errors from the one standard deviation error extracted from 1000 bootstraps (**Materials and Methods**).

### Holo-promoter dissociations occurs from RP_I_

The OS dwell times kinetics were insensitive to the holo concentration, and strictly double exponentially distributed at 150 mM KAc (**Figure 1D, Figure 2D-F, Supplementary Figure 2**), and not by a single exponential as previously reported (42). This indicates that the open complex is not formed by a single state, but suggest the existence of at least one intermediate preceding RP_O_ (8). Here, we could describe all the data in any experimental conditions with a single intermediate (RP_I_). The literature indicates that the holo very quickly isomerizes from RP_C_ to the first intermediate (rate constant *k*_2_, **Figure 2C**) in comparison to the reverse reaction (slow, rate constant *k*_−2_) and the forward reaction (8). The holo thus rapidly interconverts from RP_I_ to RP_O_. The OS dwell time distribution directly informs on the total duration of the open complex, i.e. from the promoter opening to its closing, which leads to holo dissociation. What is the kinetic pathway best describing the OS dwell times?

To answer this question, we discuss the merits of several kinetic models (**Table 1**). For each model, we determined the mathematical expressions of the rate constants describing the OS survival time distribution. For the models describing a double exponential distribution, the absolute values of the rate constants were subsequently calculated using conversion expressions as a function of the MLE fit parameters (*k*_+_, *k*_−_ and *p*_−_) (**Materials and Methods, Supplementary Information**). We first considered a model where the holo dissociates from walking back the kinetic pathway of RP_O_ formation to eventually dissociate from RP_C_ (Model 1, **Table 1**). Though this model is highly unlikely, given that *k*_2_ is much larger than *k*_−_ and *k*_+_, we calculated the resulting probability distribution function (pdf) for such model (**Supplementary Information**). We found that the pdf would be the sum of a “peaked” distribution (resembling a gamma distribution (66)) and an exponential decay, while we clearly observed a double exponential. We therefore discarded this model, and we did not consider anymore dissociation from RP_C_ in the following models. In Model 2, we consider holo dissociation only from RP_O_, which has two possible cases, i.e. starting the OS from either RP_I_ (case 1) or RP_O_ (case 2). The former, i.e. Model 2 case 1 (**Table 1**), is described by a peaked dwell time distribution (**Supplementary Information**), which is not supported by our data that consistently showed a double exponential behavior (**Supplementary Figure 2**). We therefore discarded this model. In Model 2 case 2, the open complex starts and dissociates from RP_O_ and is analytically described by a double exponential pdf (Model 2, case 2 in **Table 1, Supplementary Figure 4**). While this model is mathematically correct, it is conceptually irrational to have the open complex starting from RP_O_, and not in RP_I_ as described in the last 40 years literature (8), and we therefore discarded this model.

We introduced a third model, i.e. Model 3, where the holo may also dissociate from RP_I_ with a rate *k*_5_.

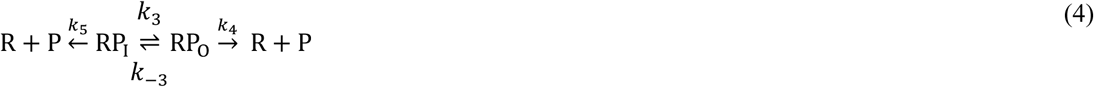

A complete mapping of the fit parameters from the double exponential pdf, i.e. *p*_+_, *k*_+_, *k*_−_, to the kinetic rates of this model (*k*_3_, *k*_−3_, *k*_4_, *k*_5_) cannot be made (**Supplementary Information**). We therefore proposed and test a set of simplifying assumptions that enable a complete mapping of the fit parameters to the underlying kinetic rates (**Table 1, Supplementary Information**). In Assumption 1, we assumed that RP_I_ and RP_O_ are in rapid equilibrium, i.e. *k*_−3_ and *k*_3_ are large in comparison to *k*_5_ and *k*_4_ (**Table 1, Supplementary Information**). Applying this model to our data resulted in negative values for *k*_4_ (**Supplementary Figure 4**), which is unphysical and therefore rejected. In Assumption 2, we defined *k*_−3_ = 0, i.e. RP_O_ is an irreversible state leading to holo dissociation (**Table 1, Supplementary Information**). While this model results in a double exponential pdf and did not produce unphysical rate values (**Supplementary Figure 4**), it implies that the only way to rescue the holo from RP_O_ is dissociation, which is in contradiction with previous studies showing that such state should be reversible without dissociation (39,43). Finally, in Assumption 3, we defined *k*_4_ = 0, i.e. the holo may enter RP_O_, but must return to RP_I_ to dissociate (**Table 1, Supplementary Information**). This model describes the OS dwell times with a double exponential pdf, without unphysical values for the rate constants (**Supplementary Figure 4**). Furthermore, it is consistent with ensemble descriptions of the holo dissociation from the λP_R_ promoter (8,67). In the following part of the study, we represented only the rate constants from Model 3 Assumption 3 (**Table 1**). We subsequently investigate how the open-complex dynamics is affected by the nature and the concentration of monovalent salts, and the temperature in the context of this model.

### Anions affect open complex dynamics and cations affect DNA twist

We first investigated how the identity of the monovalent ion affects the open complex dynamics. Previous ensemble studies showed that the DNA twist depends on the nature and the concentration of the monovalent cation (68). By affecting the DNA twist, the cation nature may impact bacterial transcription initiation kinetics (69), as the open complex dynamics is sensitive to torque (42) (**Supplementary Figure 1C-F**). In addition to the data at 150 mM potassium acetate (KAc), we investigated open complex dynamics in the presence of 150 mM of either ammonium acetate (NH_4_Ac), potassium glutamate (KGlu), sodium chloride (NaCl) or potassium chloride (KCl) (**Materials and Methods**). We used 10 nM holo in the presence of chloride anion, and 1 nM otherwise (**Figure 3**). A direct observation of the activity traces shows very long CS and very short OS dwell times in the presence of NaCl and KCl, while the opposite trend is apparent when KGlu was used. KAc and NH_4_Ac presence induced an intermediate response, i.e. CS and OS having equal durations (**Figure 3A-E**). Analyzing the dwell time distributions showed that the CS dwell times are mono-exponentially distributed in all salts (**Supplementary Figure 5)**, and *k*_*open*_ is more than three-fold larger in KGlu than in other salts (**Figure 3F**), indicating that KGlu strongly favors the open complex formation. Furthermore, we found that the OS dwell times were very short and mono-exponentially distributed in the presence of 150 mM chloride anion, while they were double-exponentially distributed in the presence of the other anions (**Figure 3G, Supplementary Figure 5**). This result suggests that the complex never reaches the stable RP_O_ in the presence of high chloride concentration and can only populate the less stable RP_I_, by which the holo dissociates from the promoter (Model 3 Assumption 3, **Table 1, Supplementary Information**). In the presence of 150 mM KAc, NH_4_Ac and KGlu, the RP_I_ state is sufficiently stable (**Figure 3I**) for the holo to visit the RP_O_ state (second exponential appearing again in the OS dwell time distributions, **Supplementary Figure 5**). We could also estimate the isomerization rate constants to (*k*_3_) and away from (*k*_−3_) the RP_O_ state (**Figure 3H, I**). The values indicate that the RP_O_ is most stable, i.e. *k*_−3_ the smallest, in KGlu whereas KAc imposes rapid dynamics between the RP_I_ and the RP_O_ states.

Could the effect we observed here resulted from a change in the DNA twist due to the change of monovalent salt? Magnetic tweezers are a well-suited technique to characterize DNA twist variation, and we therefore investigated how the DNA twist (Δtwist) varied when changing the monovalent salt from NaCl to either KCl, KGlu or NH_4_Ac. Specifically, we performed extension-rotation experiments on a 20.6 kbp coilable DNA tether (**Supplementary Figure 6, Materials and Methods**). We observed that changing the cation (sodium to potassium) in Tris-EDTA (TE) buffer induced a positive increase in twist by (135 ± 7)°/*kbp*; this cation effect was the same using either chloride or glutamate as the anion (**Supplementary Figure 6AB**). Consistently with ensemble data (68), NH_4_Ac induces even larger increase in helical twist, i.e. (331 ± 7)°/*kbp*, in comparison to NaCl. When performing the same experiments in the holo reaction buffer, which contains 5 mM MgCl_2_, we observed a similar trend, though the effect is nearly two-fold smaller (**Supplementary Figure 6B**). Our data confirms that the cation affects the DNA helical twist and the strength of this effect follows the order Na^+^<K^+^<NH_4_^+^. In contrast, our data did not show that the anion nature affects the DNA twist. We hypothesized that an increase in DNA helical twist would lead to a shorter-lived and less populated OS, as previously suggested (69). However, the observed difference in open complex dynamics (**Figure 3F-I**) is not consistent with the cation ranking for the helical twist (**Supplementary Figure 6AB**). For example, *k*_*open*_ is 6-fold larger with KCl than with NaCl, though one would expect the opposite given the helical twist ranking effect, but 3-fold smaller for NH_4_Ac than for KAc. Overall, the anion nature has a much more significant impact on open complex dynamics than the monovalent cation, and we therefore performed the following experiments using only the physiological K^+^ cation.

### Physiological concentration of glutamate favors open complex formation and stability

We next investigated how the changing concentration of chloride, acetate and glutamate affects the observed open complex dynamics at constant holo concentration. We varied the KCl concentration from 50 mM to 150 mM while using 10 nM holo in the reaction buffer. The activity traces showed shorter OS and longer CS as the KCl concentration increased (**Figure 4A**). Indeed, *k*_*open*_ decreased steadily with KCl concentration, indicating a loss in holo affinity with the promoter (**Figure 4D, Supplementary Figure 7A**). Surprisingly, we found that the OS dwell times distributions were well described by a double-exponential pdf for KCl concentration up to 100 mM. Specifically, we found that the second exponential, and therefore the RP_O_ state, was completely depopulated for KCl concentration above 100 mM (**Figure 4EF, Supplementary Figure 7A**), followed by a lower stability of RP_I_ at increasing KCl concentrations, e.g. *k*_5_ increased ~20-times when increasing KCl concentration from 50 to 150 mM (**Figure 4G**).

**Figure 4.**
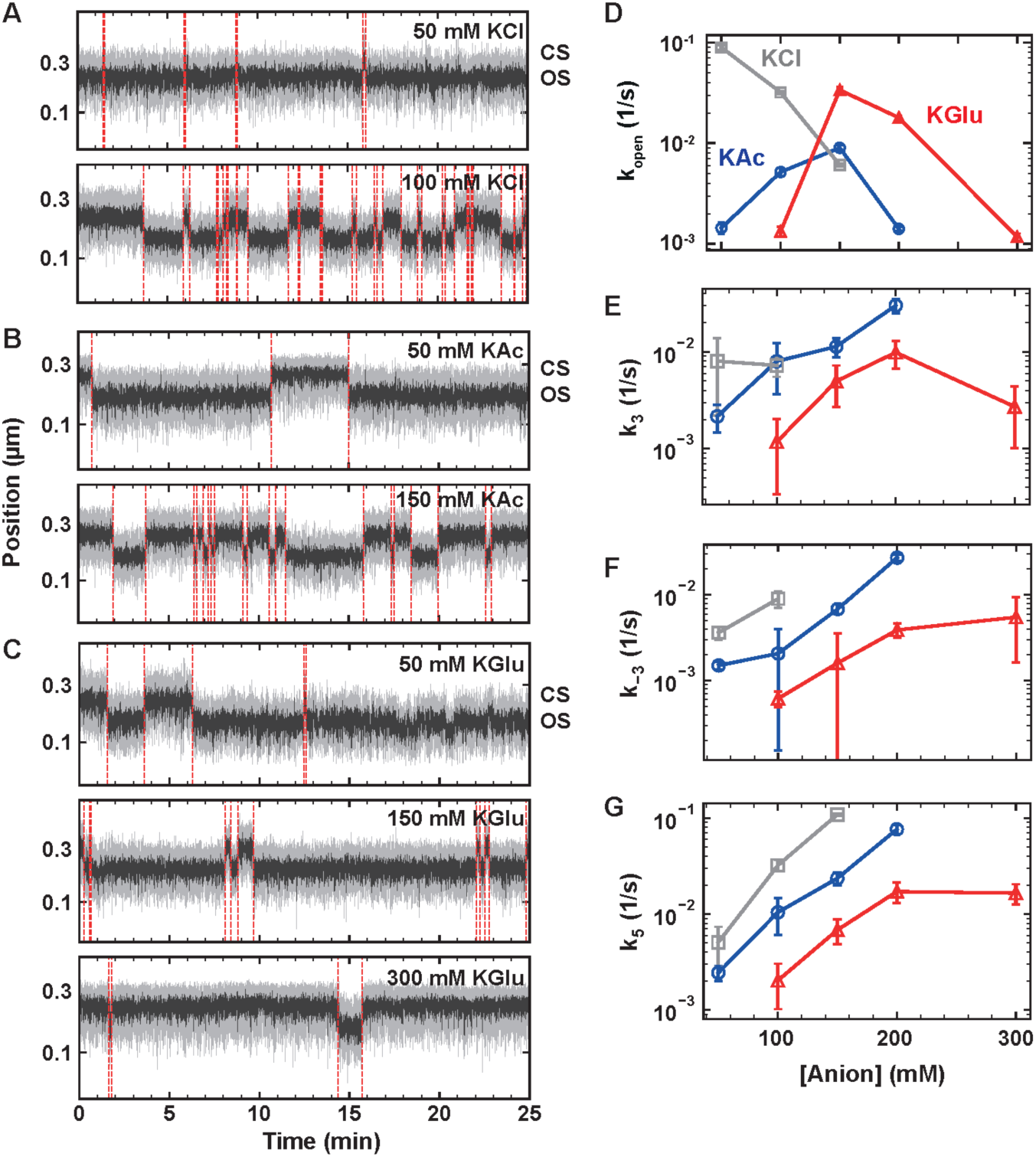
The anion type and concentration affect the holo open complex dynamics. **(A, B, C)** Traces of holo open complex dynamics using reaction buffers containing different anions at the indicated concentrations at 34°C. We used 10 nM holo in KCl, and 1 nM holo in KAc and KGlu. The red dashed lines indicate the transitions between OS and CS captured by the change-point analysis. **(D, E, F, G)** Monovalent salt concentration dependence of *k*_*open*_, *k*_3_, *k*_−3_ and *k*_5_ for KCl (grey), KAc (blue), and KGlu (red) for activity traces acquired as in (A, B, C). The solid lines connect the markers and are not fits. Error bars in (D, E, F, G) are the propagated errors from the one standard deviation error extracted from 1000 bootstraps (**Materials and Methods**)

In comparison to KCl, the holo-promoter interactions were affected in a very different way by KAc and KGlu, as *k*_*open*_ values first maximized at ~150 mM to significantly decrease at higher salt concentrations (**Figure 4BCD, Supplementary Figure 7BC**). The stability of both RP_O_ and RP_I_ decreased with KAc concentration, i.e. *k*_5_ and *k*_−3_ increased by more than one order of magnitude, while the conversion rate of RP_I_ to RP_O_, i.e. *k*_3_, surprisingly increases by more than 10-fold (**Figure 4E-G, Supplementary Figure 7BC**). While glutamate shows a destabilizing effect on the open complex similarly to acetate, the glutamate effect saturates above 200 mM concentration. We could not measure RP_O_ dynamics at KCl and KAc concentrations larger than 150 mM and 200 mM, respectively, as OS were hardly detected. RP_O_ stability in the presence of acetate is intermediate between chloride and glutamate. Interestingly, 50 mM KCl showed a faster open complex formation than 150 mM KGlu, while maintaining a comparable stability of the OS (**Figure 4D-G**). However, RP_O_ state is able to resist higher concentrations of the physiological anion glutamate than chloride, supporting the hypothesis that glutamate is an open complex stabilizer (70,71).

Having found the optimum concentration of chloride, acetate and glutamate, we investigated the open complex formation and stability as a function of holo concentration (**Figure 5**). We performed these experiments in either 100 mM KCl or 150 mM KGlu, and represent these data next to the 150 mM KAc data presented in **Figure 2**. The CS dwell times visually shortened in magnetic tweezers traces with increasing holo concentration (**Figure 5A-B**). Extracting *k*_*open*_ from the CS dwell time distributions (**Supplementary Figure 8**) and representing it as a function of holo concentration for KCl and KGlu, we observed a similar trend as for KAc and the data were well fitted by **Equation 1**, supporting that the holo rapidly dissociates from the promoter upon closing and is not recycled for the subsequent OS (**Figure 5C**). For each anion, we extracted an equilibrium association constant *K*_1_ in the 1-60 µ*M*^−1^ range and *k*_2_~1 *s*^−1^ (this value is only indicative as we could not reach saturation in holo concentration) (**Table S3**). As for KAc, the dynamics of the OS, i.e. *k*_−3_, *k*_3_ and *k*_5_, is unaffected by holo concentration in both KCl and KGlu (**Figure 5D-F**).

**Figure 5.**
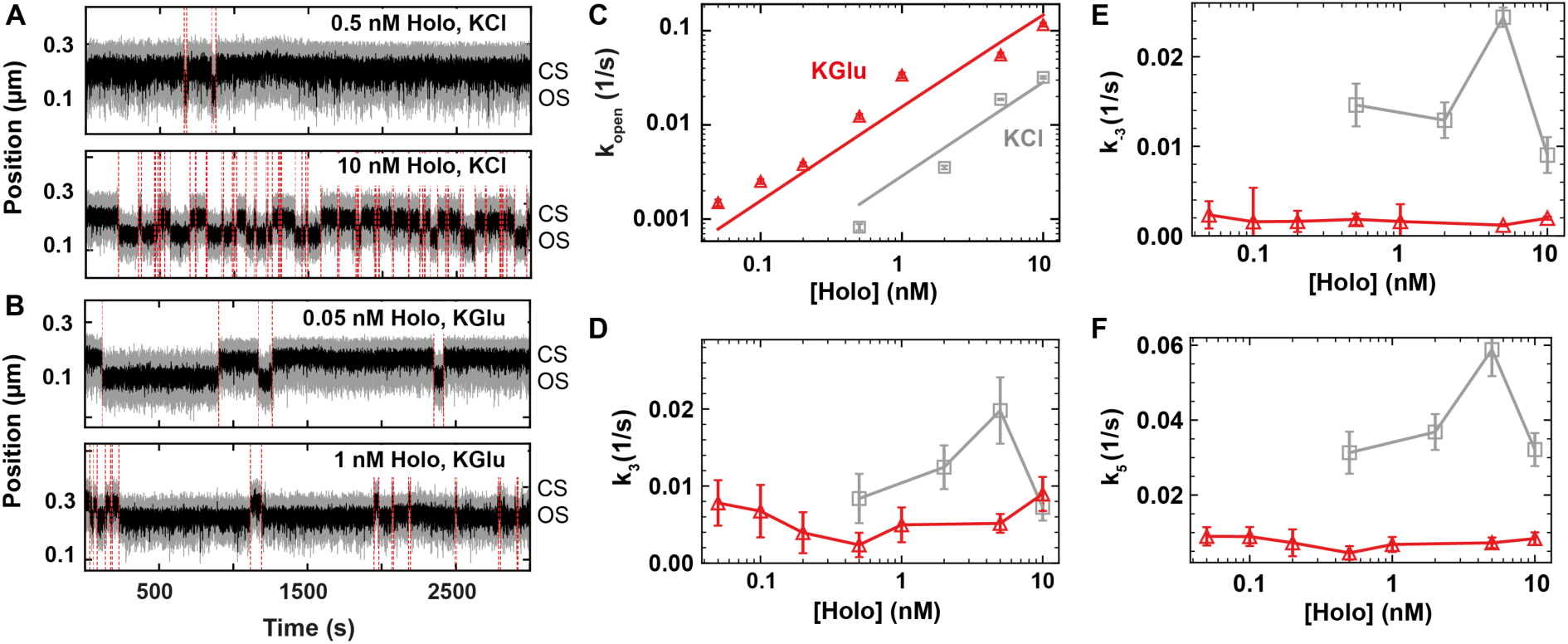
The holo releases the promoter upon transcription bubble closing. **(A, B)** Magnetic tweezers activity traces showing open complex dynamics in either (A) 100 mM KCl, or (B) 150 mM KGlu at 34°C using the indicated holo concentration. The red dashed lines indicate the transitions between OS and CS captured by the change-point analysis. **(C, D, E, F)** Holo concentration dependence of *k*_*open*_, *k*_3_, *k*_−3_ and *k*_5_ in either 100 mM KCl or 150 mM KGlu. Color code as in (C). The solid lines in (C) are fits to **Equation 1**. The solid lines in (D, E, F) connect the markers and are not fits. Error bars in (C, D, E, F) are the propagated errors from the one standard deviation error extracted from 1000 bootstraps (**Materials and Methods**)

### Open complex formation energy landscape probed by temperature-controlled magnetic tweezers

Temperature dependence of the bacterial open complex formation enables the exploration of the energy landscape of the reaction (33,72,73). We have recently developed a temperature-controlled magnetic tweezers assay (50), and we applied it to investigate how temperature affects the kinetics of the open complex dynamics in real-time. Because KGlu induces extremely stable OS, we performed this study in 150 mM KAc and 5 nM holo to maximize the statistics of the open complex dynamics as a function of temperature (**Figure 6A**). Nonetheless, we expect our results to be conserved for KGlu, as the open complex dynamics shows a similar trend in either acetate or glutamate (**Figure 4**).

**Figure 6.**
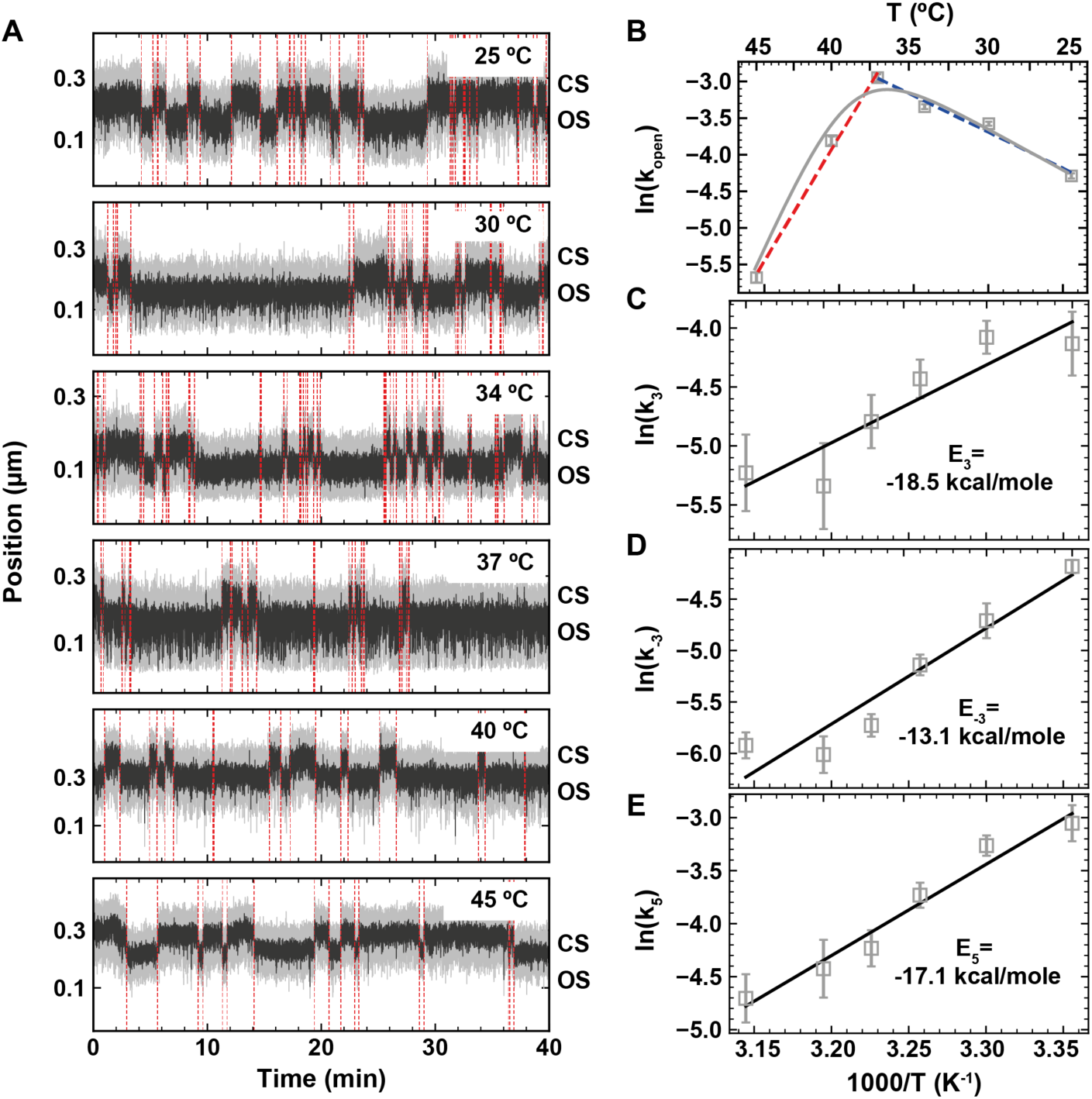
Effect of temperature on the holo open complex dynamics. All experiments were performed in 150 mM KAc and 5 nM holo. **(A)** Magnetic tweezers activity traces showing open complex dynamics at the temperature indicated in the plots. The red dashed lines indicate the transitions between OS and CS captured by the change-point analysis. **(B)** Arrhenius plot of *k*_*open*_ where the solid line is a fit to the data using **Equation 3**. Blue and red dashed lines in (B) are fits using **Equation 2** restricted to *k*_*open*_ extracted for temperatures either below or above 37°C, respectively. The energy landscape of the open state formation is represented within the plot. **(C, D, E)** Arrhenius plot of *k*_3_, *k*_−3_ and *k*_5_. Solid lines are fit to the data using **Equation 2**. Error bars in B, C, D, E are the propagated errors from the one standard deviation error extracted from 1000 bootstraps (**Materials and Methods**)

We showed here that the CS dwell times distribution was strictly mono-exponential described by an exit rate *k*_*open*_ that is strongly holo concentration dependent (**Supplementary Figure 9**). Therefore, if the holo was degraded/denatured during the course of the experiment (several hours) at the elevated temperature, we would have expected the CS dwell times distribution to not be accurately fitted by a mono-exponential, but a probability distribution function representative of the holo concentration decrease over time, i.e. a multi-exponential. The observed CS dwell time distributions at all temperatures indicate that the holo remained functional for the entire duration of the experiment. The MLE fits revealed that *k*_*open*_ increased by ~4-fold when temperature was increased from 25 °C to 37 °C, and subsequently decreased by ~13-fold when the temperature further increased from 37 °C to 45 °C (**Supplementary Figure 9, Figure 6B**). *k*_*open*_ cannot be fitted by a simple Arrhenius equation, as the Arrhenius plot does not appear curvilinear (**Figure 6B**) (74). This behavior is similar to what was previously described for the temperature dependence of fibrinopeptide release by thrombin (75). The transition with the lowest activation energy, i.e. from RP_C_ to RP_I_, dominates the reaction at low temperature, while the transition with the highest activation energy dominates at high temperature, i.e. holo dissociation from the promoter. Such behavior was not described in the previous investigations of the temperature dependence of the open complex formation with strong promoters, likely because holo dissociation never dominated (33,72,73). Using **Equation 3** (**Materials and Methods**), we extracted the activation energy of the transition from RP_C_ to RP_I_, i.e. Δ*E*_2_ = (22 ± 6) *kcal/mol*, and the energy difference between the unbound state R+P and RP_C_, i.e. Δ*E*_*diff*_ = (107 ± 12) *kcal/mol* (**Figure 6B**). Our evaluation of Δ*E*_2_ is in agreement with previous estimation made using the *lacUV5* promoter (33,73).

From the OS dwell time distributions (**Supplementary Figure 9**), we extracted the temperature dependence of *k*_3_, *k*_−3_ and *k*_5_. Their respective Arrhenius plot were well described by **Equation 2**, and we extracted the activation energies *E*_3_ = (−18.5 ± 2.9) *kcal* · *mol*^−1^, *E*_−3_ = (−13.1 ± 3.1) *kcal* · *mol*^−1^ and *E*_5_ = (−17.1 ± 1.7) *kcal* · *mol*^−1^, respectively (**Figure 6C-E**). Interestingly, all these activation energies are negative, which indicates the presence of intermediate states trapping the complex in metastable states on the pathway to dissociation. While for *lacUV5* and *λP*_*R*_, negative *E*_−3_ and *E*_5_ were already observed, we also note that *E*_3_ is negative, indicating the existence of another intermediate between RP_I_ and RP_O_, as previously suggested for *λP*_*R*_ promoter, i.e. I_3_ (35,76).

## Discussion

In the presented study, we have investigated the bacterial open complex formation and dissociation on a consensus *lacUV5* (*lacCONS*) promoter using high throughput magnetic tweezers. To this end, we have studied the impact of the nature and concentration of the monovalent salt, holo concentration and temperature on the kinetics of formation and dissociation. While some of these aspects have been investigated in ensemble studies with *λ*P_R_ and *lacUV5* promoters, such investigations have never been performed at the single molecule level. Furthermore, we show here that the choice of the monovalent salt may have dramatic consequences on the kinetics observed at the single molecule level. Seminal work by Strick and colleagues using magnetic tweezers showed that the holo specifically binds at the promoter to form an RP_C_ and transits directly towards a stable RP_O_, the lifetime of which varies as a function of the promoter sequence, the applied torque and the supercoiling sign (42). Interestingly, they reported no intermediate between RP_C_ and RP_O_, and a single dissociation rate constant, though ensemble studies already reported at least one intermediate (8). Furthermore, a recent biochemical study has characterized several closed-promoter intermediates preceding RP_I_, which originate from promoter bending and conformational rearrangement of the RNAP clamp to position the promoter towards opening (26), supporting recent cryoEM studies (25,77). A recent single-molecule FRET study has showed that the holo explores an RP_I_ state, either transiently or permanently, in addition to the fully open RP_O_ state (43). One of the differences between the magnetic tweezers and the single molecule FRET studies was the nature of the monovalent salt: the former used NaCl, while the latter used KGlu. Record and colleagues have shown that the physiologically relevant glutamate has a stabilizing effect on protein folding and RP_O_ formation over chloride, and specifically interacts with the holo to drive major conformational changes from RP_I_ (called I_2_ in *λ*P_R_ studies) to RP_O_ (71). These experiments were performed at rather high monovalent salt concentration, i.e. from 150 mM to 545 mM, and did not investigate lower, and more physiological, salt concentrations. We have filled this gap by investigating a range of monovalent salts from 50 mM to 300 mM. We show here that open complex dynamics are mainly affected by the type of the anions, i.e. Cl^-^, Glu^-^ and Ac^-^. In agreement with previous studies (70,78), our direct observation of RP_O_ formation using a torsionally constrained DNA molecule showed no effect of the monovalent cation, i.e. K^+^, Na^+^ and NH_4_^+^(**Figure 3G-I**), despite the increase in the DNA helical twist induced by potassium and ammonium in comparison to sodium (**Supplementary Figure 6**). Though the rate of open complex formation was similar at 50 mM KCl and 150 mM KGlu, it decreased exponentially with KCl concentration (**Figure 4D**), until no activity was detected above 150 mM. This result suggests that increasing chloride concentration screens the holo Coulombic interactions with the promoter, and consequently decreases the equilibrium association constant *K*_1_. KGlu and KAc impacts the open complex formation in two different regimes, up to ~150 mM, KGlu increases attractive interactions between the promoter and the holo, to eventually screen these interactions by Coulombic effect at higher concentrations.

We studied the dissociation kinetics of the holo from *lacCONS* promoter to determine the nature of RP_O_ and RP_I_. Kinetic modeling clearly supports a description where RP_O_ is a stable but still slowly reversible state and holo dissociation occurs only via RP_I_ (Model 3 Assumption 3, **Table 1**). This is in agreement with ensemble studies inducing dissociation with high salt upshift (67). How does anion concentration affect open complex stability? The dissociation rate constant *k*_5_, i.e. from RP_I_ to R+P, increased exponentially with glutamate and acetate concentration, and even more so for chloride, up to the point that the holo did not enter RP_O_ state at >100 mM chloride. The strong dependence of RP_I_ dissociation on chloride could explain why a previous magnetic tweezers study reported only a single rate constant for RP_O_ dissociation, as if the entire OS population was in the RP_I_ state (42). Interestingly, a follow up study by the same group on initial transcription and promoter escape showed RNA synthesis activity in the same ionic condition, suggesting that RP_I_ may be the catalytically competent state (79), as recently reported by Record and colleagues (76). We could not observe a difference in transcription bubble size between RP_I_ and RP_O_, indicating that this transition does not increase the bubble size, but rather rearranges the melted DNA strands inside the holo (70) and/or involves conformation change in the holo. These rearrangements tighten the holo–promoter interactions effectively preventing the dissociation from RP_O_ state. In the RP_I_ state, in contrast, these interactions remain weak enough to allow their stochastic disruption and thus direct dissociation from the RP_I_ state without full reversal to the state preceding the open complex, i.e., RP_C_. Furthermore, acetate shows an intermediate effect on the kinetics of *k*_3_, *k*_−3_ and *k*_5_, following the Hofmeister series ranking of acetate between chloride and glutamate, and being also consistent with the idea that glutamate favors open complex formation via non-Coulombic interactions (70).

By investigating the temperature dependence of the RP_O_ dynamics, we extracted the activation energies of the transitions between the different states. We showed that, in our experimental conditions, i.e. positively supercoiled DNA and 150 mM KAc, a simple Arrhenius equation could not describe the temperature dependent open complex formation kinetics (**Figure 6B**). We thus proposed a model, in which the transition from RP_C_ to RP_I_ (*k*_2_ in **Figure 2C**) dominates the reaction at low temperature, while RP_C_ to holo dissociation (*k*_−1_ in **Figure 2C**) dominates the reaction at high temperature. As previously observed (33,76), the activation energies of the transition described by the reaction rate constants *k*_3_, *k*_−3_ and *k*_5_ are negative, suggesting that the transitions from RP_O_ to RP_I_ (and vice versa), and from RP_I_ to R+P do not occur in a single step, but rather involve intermediates that trap the complex during the net dissociation reaction, though not being kinetically significant in the net forward reaction of RP_O_ formation.

Our study expands the understanding on how monovalent salts and temperature affect protein-nucleic acids interactions, and will therefore be of use to single molecule biophysicists. We provide the community a detailed protocol to establish a robust lipid bilayer passivation for single molecule assays, as well as a complete pipeline for data analysis using a custom Python routine. Furthermore, our single molecule study of bacterial RNA polymerase open complex dynamics corroborates the existence of open-state intermediates, and further expands our understanding of the interactions leading to a stable RNAP open complex. Future studies will focus on the direct observation of these intermediates using high resolution magnetic tweezers combined with single molecule FRET to reveal the complete cycle of open complex formation and dissociation.

## Supporting information

Supplementary Information for Bera, America et al.

## Data availability

The data of this study are available from the authors upon reasonable request.

## Supplementary Data

Supplementary Data are available at NAR online.

## Acknowledgements

DD was supported by the Interdisciplinary Center for Clinical Research (IZKF) at the University Hospital of the University of Erlangen-Nuremberg, and the German Research Foundation grant DFG-DU-1872/3-1, DFG-DU-1872/4-1, DFG-DU-1872/5-1 and BaSyC – Building a Synthetic Cell” Gravitation grant (024.003.019) of the Netherlands Ministry of Education, Culture and Science (OCW) and the Netherlands Organisation for Scientific Research (NWO). AMM was supported by the Academy of Finland (grant numbers 307775, 314100 and 335377). DD would like to thank OICE for hosting his research group. DD thanks Jan Lipfert for insightful discussions.

## Author contributions

AMM and DD designed the research. DD supervised the research. SCB performed the single molecule magnetic tweezers experiments, analyzed and represented the data. MSe wrote the initial analysis step finder program. EO developed the lipid bilayer functionalization strategy. JC provided computer analysis routine and support for the computer set-up Labview interface. AMM and SM provided the *E. coli* holoenzyme complex. FSP made the nucleic acid constructs. MD and PAA derived the mathematical expression for the kinetic model. MSp performed initial experiments. All authors have contributed in the discussion and interpretation of the data. SCB, AMM and DD wrote the article. All authors have edited the article.

