## Supplementary Information for Bera, America et al. for "Quantitative parameters of bacterial RNA polymerase open-complex formation, stabilization and disruption on a consensus promoter"

**This Supplementary Information contains**

Supplementary Notes 1 to 2

Supplementary Figures 1 to 9

Table S1 to S3

**Supplementary notes 1: Kinetic description of double exponential OS dwell time distributions**

In this section, we discuss how we evaluated which kinetic model best describes the open state (OS) dwell time distribution, or in other words, the OS disassembly time. First note that the unbinding distribution for a kinetic model can in general be obtained by evaluating the first passage time distributions of all the paths that result in unbinding. If we assume the specific transition from $x$ to $y$ occurs with rate $k_{x\to y}$, when the total rate of all transitions starting from state $x$ is $k_{\mathrm{tot,x}}$, we can write the first passage time distribution for the transition as

$$p_{x\to y}(t) = p_{x\to y} k_{\mathrm{tot,x}} e^{-k_{\mathrm{tot,x}} t} = k_{x\to y} e^{-k_{\mathrm{tot,x}} t}.$$

Here we have used that the (splitting) probability for making the transition to $x$ before any of the other transitions is

$$p_{x\to y} = \frac{k_{x\to y}}{k_{\mathrm{tot,x}}}.$$

The first passage time distribution of a path through multiple states can be found as the convolution of the first passage time distributions between the successive states. To enable to combine successive transitions by simply multiplying probability densities (rather than performing convolutions), we move to Laplace space, and write the general probability distribution as

$$\phi_x(s) = \int_0^\infty e^{-st} p_x(t) \mathrm{d}t = \frac{k_x}{s + k_{\mathrm{tot}}}$$

**Model 1: open complex starts from $RP_I$ and dissociates from $RP_C$**

We first evaluate the most straight-forward model for OS dissociation, i.e. the holo enters the OS from $RP_I$ to reach $RP_O$, and walks back the kinetic pathway to dissociate from $RP_C$, such as

$$\mathrm{R} + \mathrm{P} \underset{k_{-1}}{\longleftarrow} \mathrm{RP_C} \underset{k_{-2}}{\overset{k_2}{\rightleftharpoons}} \mathrm{RP_I} \underset{k_{-3}}{\overset{k_3}{\rightleftharpoons}} \mathrm{RP_O} \qquad \text{(S1)}$$

The system converts from the $RP_I$ to $RP_C$ or $RP_O$ with rate $k_{-2}$ or $k_3$ respectively, and can get repeatedly reabsorbed back into $RP_I$ with rate $k_2$ or $k_{-3}$ before the holo dissociates from $RP_C$ with rate $k_{-1}$. Of note, this model can be interpreted as an extension of **Model 2, Assumption 3**, where $k_4$ is subdivided into $k_2$, $k_{-2}$ and $k_{-1}$.

Using this kinetic model, we have the following transition-time distributions

$\phi_{-1} = \frac{k_{-1}}{k_{-1}+k_2+s}$ , $\phi_2 = \frac{k_2}{k_{-1}+k_2+s}$, $\phi_{-2} = \frac{k_{-2}}{k_3+k_{-2}+s}$

$\phi_3 = \frac{k_3}{k_3+k_{-2}+s}$, $\phi_{-3} = \frac{k_{-3}}{k_{-3}+s}$

By summing transition-time distributions of escape paths, enumerated by the number $n$ of absorptions into $RP_O$ before escaping, we can write down the full escape-time distribution. Starting from $RP_I$, the escape-time distribution is

$$\Psi_{RP_I \to R+P}(s) = \phi_{-1}\phi_{-2}\sum_{m_1}^{\infty}\sum_{m_2}^{\infty}\frac{(m_1+m_2)!}{m_1!\,m_2!}(\phi_2\phi_{-2})^{m_1}(\phi_3\phi_{-3})^{m_2}$$

Defining $m = m_1 + m_2$, we have

$$\Psi_{RP_I \to R+P}(s) = \phi_{-1}\phi_{-2}\sum_{m}^{\infty}\sum_{m_1}^{m}\frac{m!}{m_1!\,(m-m_1)!}(\phi_2\phi_{-2})^{m}(\phi_3\phi_{-3})^{m-m_1}$$

Using the binomial theorem, we can write

$$\Psi_{RP_I \to R+P}(s) = \phi_{-1}\phi_{-2}\sum_{m}^{\infty}(\phi_2\phi_{-2}+\phi_3\phi_{-3})^{m} = \frac{\phi_{-1}\phi_{-2}}{1-\phi_2\phi_{-2}-\phi_3\phi_{-3}}$$

$$= \frac{k_1k_{-2}(k_{-3}+s)}{(k_{-1}+k_2+s)(k_3+k_{-2}+s)(k_{-3}+s)-k_2k_{-2}(k_{-3}+s)-k_3k_{-3}(k_{-1}+k_2+s)}$$

$$= \frac{k_{-1}k_{-2}(k_{-3}+s)}{(s+s_1)(s+s_2)(s+s_3)}$$

Where $s_1$, $s_2$ and $s_3$ are defined as minus the roots of the denominator. As we have three roots, we can write $\Psi_{RP_I \to R+P}$ as the sum of three Laplace transformed exponentials using partial fraction decomposition, such as

$$\Psi_{RP_I \to R+P}(s) = \frac{A_1}{s+s_1}+\frac{A_2}{s+s_2}+\frac{A_3}{s+s_3}$$

$$= \frac{A_1(s+s_2)(s+s_3)+A_2(s+s_1)(s+s_3)+A_3(s+s_1)(s+s_2)}{(s+s_1)(s+s_2)(s+s_3)}$$

$$= \frac{s^2(A_1+A_2+A_3)+s[A_1(s_2+s_3)+A_2(s_1+s_3)+A_3(s_1+s_2)]+A_3s_1s_2+A_1s_2s_3+A_2s_1s_3}{(s+s_1)(s+s_2)(s+s_3)}$$

Comparison of the numerators gives us for the following relations

$$A_1+A_2+A_3 = 0$$

$$A_1(s_2 + s_3) + A_2(s_1 + s_3) + A_3(s_1 + s_2) = k_{-1}k_{-2}$$

$$A_3 s_1 s_2 + A_1 s_2 s_3 + A_2 s_1 s_3 = k_{-1}k_{-2}k_{-3}$$

Considering the probability density function must be normalized in the time domain, we can write it in the form

$$\mathrm{P}_{RP_I \to R+P}(t) = p_1 s_1 e^{-s_1 t} + p_2 s_2 e^{-s_2 t} + (1 - p_1 - p_2) s_3 e^{-s_3 t}$$

Note that in this notation $A_1 = p_1 s_1$, $A_2 = p_2 s_2$ and $A_3 = (1 - p_1 - p_2)s_3$, so we can rewrite the relations from the numerator to

$$p_1 s_1 + p_2 s_2 + (1 - p_1 - p_2)s_3 = 0 \qquad \text{(S2)}$$

$$p_1 s_1(s_2 + s_3) + p_2 s_2(s_1 + s_3) + (1 - p_1 - p_2)s_3(s_1 + s_2) = k_{-1}k_{-2} \qquad \text{(S3)}$$

$$s_1 s_2 s_3 = k_{-1}k_{-2}k_{-3} \qquad \text{(S4)}$$

From **Equations S2** and **S3**, we get

$$p_1 = \frac{s_2 s_3 - k_{-1}k_{-2}}{(s_3 - s_1)(s_2 - s_1)} \qquad \text{(S5)}$$

$$p_2 = -\frac{s_1 s_3 - k_{-1}k_{-2}}{(s_3 - s_2)(s_2 - s_1)} \qquad \text{(S6)}$$

$$p_3 = 1 - p_1 - p_2 = \frac{s_1 s_2 - k_{-1}k_{-2}}{(s_3 - s_2)(s_3 - s_1)} \qquad \text{(S7)}$$

We now need to evaluate the signs of the exponential weights to determine the nature of the distribution, considering that the kinetic rates must be positive.

First note that we only consider distributions with real and positive exponential rates (resulting in exponential decay) such that we can choose $s_3 > s_2 > s_1 > 0$. Starting from **Equations S2** and **S3**, we obtain

$$k_{-1}k_{-2} = -p_1(s_3 - s_1)(s_2 - s_1) + s_2 s_3$$

Considering that $k_{-1}, k_{-2} > 0$ we find the condition

$$p_1 > \frac{s_2 s_3}{(s_3 - s_1)(s_2 - s_1)} > 0 \qquad \text{(S8)}$$

Of note, we have $\frac{s_2}{(s_2 - s_1)} > 1$ and $\frac{s_3}{(s_3 - s_1)} > 1$ because $s_3 > s_2 > s_1 > 0$. We can therefore rewrite **Equation S8** such as

$p_1 > \frac{s_3}{s_3 - s_1} > 1$ and $p_1 > \frac{s_2}{s_2 - s_1} > 1$

The numerator of **Equation S6** must verify

$k_{-1}k_{-2} - s_1 s_3 = (s_2 - s_1)[-p_1(s_3 - s_1) + s_3] < 0$ and the denominator is positive, therefore $p_2 < 0$.

Furthermore, the numerator of **Equation S7** must verify

$s_1 s_2 - k_{-1}k_{-2} = (s_3 - s_1)[p_1(s_2 - s_1) - s_2] > 0$, the denominator is positive and therefore $p_3 > 0$.

In conclusion, we have two exponentials with a positive weight and one with a negative rate. The global distribution is therefore the combination of a peaked and an exponential distribution.

**Model 2: Dissociation from RP$_O$**

As dissociation from RP$_C$ (Model 1) fails at describing the double exponential distribution of the OS dwell times, we propose another model, where the holo dissociates from RP$_O$.

$$\mathrm{RP_I} \underset{k_{-3}}{\overset{k_3}{\rightleftharpoons}} \mathrm{RP_O} \overset{k_4}{\rightarrow} \mathrm{R} + \mathrm{P} \tag{S9}$$

The system converts from the RP$_I$ to RP$_O$ with rate $k_3$, and can get temporarily and repeatedly reabsorbed back into RP$_I$ with rate $k_{-3}$, before the holo dissociates from RP$_O$ with rate $k_4$. In the following section, we calculate the escape time distribution from the double bound states system for two cases, i.e. the OS starts in either RP$_O$ or RP$_I$.

The particular transition-time distributions for **Equation S9** are

$$\phi_4(s) = \frac{k_4}{s + k_4 + k_{-3}}, \qquad \phi_3(s) = \frac{k_3}{s + k_3}, \qquad \phi_{-3}(s) = \frac{k_{-3}}{s + k_4 + k_{-3}}.$$

By summing transition-time distributions of escape paths, enumerated by the number $n$ of absorptions into RP$_O$ before escaping, we can write down the full escape-time distribution, which differs depending on the starting point of the OS.

**Model 2, Case 1: the OS starts in RP$_O$**

Here, the OS starts in RP$_O$, and therefore we have

$$\Psi_{RP_O\to R+P}(s) = \phi_4(s)\sum_{n=0}^{\infty}\left(\phi_3(s)\phi_{-3}(s)\right)^n = \frac{\phi_4(s)}{1-\phi_{-3}(s)\phi_3(s)} = \frac{k_4(s+k_3)}{(s+k_3)(s+k_4+k_{-3})-k_{-3}k_3}$$

This escape time distribution is readily converted back from Laplace to real space as

$$P_{RP_O\to R+P}(t) = p_+k_+e^{-k_+t} + (1-p_+)k_-e^{-k_-t}.$$

With $p_{i,+} = \frac{A_{i,+}}{k_+}$ and $1-p_+ = \frac{A_{i,-}}{k_-}$.

We have the two effective rates

$$k_\pm = \frac{1}{2}(k \pm \Delta k), \qquad k = k_{-3}+k_3+k_4, \qquad \Delta k = \sqrt{k^2-4k_3k_4}$$

By definition, we have $k_\pm$

$$k_+ + k_- = k_{-3}+k_3+k_4$$

If we now find the roots of the denominator to factorize it and use partial-fraction decomposition we find that $\Psi_{RP_O\to R+P}(s)$ can be written in the form of two Laplace transformed exponential.

$$\Psi_{RP_O\to R+P}(s) = \frac{k_4(s+k_3)}{(s+k_3)(s+k_4+k_{-3})-k_{-3}k_3} = \frac{s(A_{O,+}+A_{O,-})+A_{O,+}k_-+A_{O,-}k_+}{(s+k_+)(s+k_-)}$$
$$= \frac{A_{O,+}}{s+k_+} + \frac{A_{O,-}}{s+k_-}$$

The numerator of $\Psi_{RP_O\to R+P}$ must follow

$A_{O,+}+A_{O,-} = k_4$ and $A_{O,+}k_- + A_{O,-}k_+ = k_3k_4$

Which can also be written as

$k_4 = p_+k_+ + (1-p_+)k_-$ and $k_+k_- = k_3k_4$

From which can be derived

$p_{O,+} = \frac{k_4-k_-}{k_+-k_-}$ and $p_{O,-} = 1-p_{O,+} = \frac{k_+-k_4}{k_+-k_-}$

As we have per definition $k_+ > k_-$ ; $k_4 = p_+k_+ + (1-p_+)k_-$ and the kinetic rates should be positive, we can derive that $k_+ > k_4 > k_-$ and therefore the two exponential weights $p_{O,+/-}$ must be both positive. The kinetic rates for this model can be found from the double exponential fit parameters with the following conversion relations.

$k_4 = p_+ k_+ + (1 - p_+) k_-$

$k_3 = \frac{k_+ k_-}{k_4}$

$k_{-3} = \frac{(k_+ - k_4)(k_4 - k_-)}{k_4}$

This model is however inconsistent with the start of the OS in $RP_I$, as described in the literature, and can therefore be discarded.

**Model 2, Case 2: the OS starts in $RP_I$**

Starting in OS from $RP_I$, we need to add an extra transition from $RP_I$ to $RP_O$ state

$$\Psi_{RP_I \to R+P}(s) = \phi_3(s)\Psi_{RP_O \to R+P}(s) = \frac{\phi_3(s)\phi_4(s)}{1 - \phi_{-3}(s)\phi_3(s)} = \frac{k_3 k_4}{(s + k_3)(s + k_4 + k_{-3}) - k_{-3} k_3}$$
$$= \frac{s(A_{I,+} + A_{I,-}) + A_{I,+} k_- + A_{I,-} k_+}{(s + k_+)(s + k_-)} = \frac{A_{I,+}}{s + k_+} + \frac{A_{I,-}}{s + k_-}.$$

Similarly to Case 1, this escape time distributions is readily converted back from Laplace to real space as

$$P_{RP_I \to R+P}(t) = p_+ k_+ e^{-k_+ t} + (1 - p_+) k_- e^{-k_- t}.$$

With $p_{i,+} = \frac{A_{i,+}}{k_+}$ and $1 - p_+ = \frac{A_{i,-}}{k_-}$.

We have the two effective rates

$$k_\pm = \frac{1}{2}(k \pm \Delta k), \qquad k = k_{-3} + k_3 + k_4, \qquad \Delta k = \sqrt{k^2 - 4k_3 k_4}$$

Of note, these effective rates are independent from the starting position, i.e. either $RP_O$ or $RP_I$.

From the numerator of $\Psi_{RP_I \to R+P}$, we have

$p_+ k_+ + (1 - p_+) k_- = 0$ and $k_+ k_- = k_3 k_4$

From which can be derived

$p_{I,+} = -\frac{k_-}{k_+ - k_-} < 0, \qquad p_{I,-} = 1 - p_{O,+} = \frac{k_+}{k_+ - k_-} \geq 0$

In this case we find that one of the exponential weights is negative, which leads to a peaked distribution where a minimum of two successive steps is needed before holo dissociation (1,2). We clearly observe a double exponential distribution in **Figure 1D**, and therefore Model 2, Case 2 cannot describe the data.

**Model 3: open complex starts from $RP_I$, dissociation from $RP_O$ and $RP_I$ considered**

After concluding that **Model 1 and 2** are not suitable to describe the OS dwell time distribution, we propose a third model, which allows the holo to dissociated from both $RP_I$ and $RP_O$, while the OS starts in $RP_I$ (**Table 1**).

$$\mathrm{R+P} \overset{k_5}{\leftarrow} \mathrm{RP_I} \underset{k_{-3}}{\overset{k_3}{\rightleftharpoons}} \mathrm{RP_O} \overset{k_4}{\rightarrow} \mathrm{R+P} \qquad \text{(S10)}$$

Using the formalism described above, the particular transition-time distributions for the new model (**Equation S10**) are

$$\phi_5 = \frac{k_5}{k_5+k_3+s}, \qquad \phi_3 = \frac{k_3}{k_5+k_3+s}, \qquad \phi_{-3} = \frac{k_{-3}}{k_4+k_{-3}+s}, \qquad \phi_4 = \frac{k_4}{k_4+k_{-3}+s}$$

In this model we start from $RP_I$ and there are escape-routes with we have that the path to unbinding is

$$\Psi_{RP_I \to R+P}(s) = (\phi_5 + \phi_3\phi_4)\sum_{n=0}^{\infty}(\phi_3\phi_{-3})^n = \frac{\phi_5 + \phi_3\phi_4}{1-\phi_3\phi_{-3}} = \frac{k_5(k_4+k_{-3}+s)+k_3k_4}{(k_4+k_{-3}+s)(k_5+k_3+s)-k_3k_{-3}}$$
$$= \frac{k_5k_4 + k_5k_{-3} + k_3k_4 + k_5 s}{(s+k_+)(s+k_-)}$$

In the last step, we factorized the denominator with the roots

$$k_\pm = \frac{1}{2}(k \pm \Delta k), \qquad k = k_{-3} + k_3 + k_4 + k_5 \quad ,$$

$$\Delta k = \sqrt{k^2 - 4[(k_5+k_3)(k_4+k_{-3}) - k_3k_{-3}]} = \sqrt{k^2 - 4[k_5k_4 + k_5k_{-3} + k_3k_4]}$$

Using partial-fraction decomposition, we find that we can write $\Psi_{RP_I \to R+P}$ in the form

$$\Psi_{RP_I \to R+P} = \frac{k_5k_4 + k_5k_{-3} + k_3k_4 + k_5 s}{(s+k_+)(s+k_-)} = \frac{s(A_+ + A_-) + A_+k_- + A_-k_+}{(s+k_+)(s+k_-)} = \frac{A_+}{s+k_+} + \frac{A_-}{s+k_-}$$

Where $A_+$ and $A_-$ are a combination of the kinetic rates. Note that if we transform this form of $\Psi_{RP_I \to R+P}$ back from Laplace space to real space, we get a double exponentials probability distribution function

$$P_{RP_I \to R+P}(t) = A_+ e^{-k_+ t} + A_- e^{-k_- t} = p_+ k_+ e^{-k_+ t} + (1 - p_+) k_- e^{-k_- t}$$

To get the coefficients $A_+$ and $A_-$, note from the numerator of $\Psi_{RP_I \to R+P}$ that

$$A_+ + A_- = k_5 \quad \text{and} \quad A_+ k_- + A_- k_+ = k_5 k_4 + k_5 k_{-3} + k_3 k_4 \qquad \text{(S11, S12)}$$

From the numerator of $\Psi_{RP_I \to R+P}$ we find

$$k_+ k_- = k_5 k_4 + k_5 k_{-3} + k_3 k_4 \qquad k_+ + k_- = k_{-3} + k_3 + k_4 + k_5 \qquad \text{(S13, S14)}$$

Therefore, we can write

$$p_+ = \frac{A_+}{k_+} = \frac{k_5 - k_-}{k_+ - k_-} \quad \text{and} \quad p_- = 1 - p_+ = \frac{k_+ - k_5}{k_+ - k_-}$$

To extract the values of the kinetic rates in the model from the double exponential fits, we need expressions for the kinetic rates in terms of the fitting parameters.

First note that

$$k_5 = A_+ + A_- = p_+ k_+ + (1 - p_+) k_- \qquad \text{(S15)}$$

Secondly, consider that from **Equation S13** and **S14** we obtain

$$(k_+ + k_-)(k_3 + k_5) - k_+ k_- = {k_3}^2 + 2k_3 k_5 + {k_5}^2 + k_3 k_{-3}$$

Rewriting this relation gives

$$k_{-3} = \frac{-(k_3 + k_5)^2 + (k_+ + k_-)(k_3 + k_5) - k_+ k_-}{k_3} \qquad \text{(S16)}$$

$$= \frac{(k_+ - (k_3 + k_5))((k_3 + k_5) - k_-)}{k_3}$$

and using $k_+ + k_- = k_{-3} + k_3 + k_4 + k_5$ we get

$$k_4 = \frac{k_3 k_5 + {k_5}^2 - k_5(k_+ + k_-) + k_+ k_-}{k_3} = k_5 - \frac{(k_+ - k_5)(k_5 - k_-)}{k_3} \qquad \text{(S17)}$$

Now we have derived solutions for $k_5$, $k_{-3}$ and $k_4$ in terms of $p_+$, $k_+$, $k_-$, $k_3$. We could however not retrieve a complete set of conversion relations for fit parameters $p_+$, $k_+$, $k_-$ to kinetic parameters $k_3$, $k_{-3}$, $k_4$, $k_5$ without making an assumption. This is simply because a direct mapping of three parameters ($p_+$, $k_+$, $k_-$) to four parameters ($k_3$, $k_{-3}$, $k_4$, $k_5$) cannot be made.

**Model 3, Assumption 1: $RP_I$ and $RP_O$ in rapid equilibrium**

If we assume that the rates between states $RP_I$ and $RP_O$ ($k_3$ and $k_{-3}$) are much higher than the rates to unbind from either of the two states ($k_4$ and $k_5$), we can assume that the transitions between the states are equilibrated and the detailed balance holds between the two states

$$p_{RP_I} k_3 = p_{RP_O} k_{-3}, \qquad p_{RP_I} + p_{RP_O} = 1,$$

where $p_{RP_I}$ and $p_{RP_O}$ are the equilibrated fractions in the $RP_I$ and $RP_O$ state respectively. In this case, we have a double exponential unbinding distribution with $p_+ = p_{RP_I}$ and $1 - p_+ = p_{RP_O}$. Thus, we have $k_{-3} = \frac{p_{RP_I}}{p_{RP_O}} k_3 = \frac{p_+}{1-p_+} k_3$. Substitution of this relation in the general expression for $k_{-3}$ (**Equation S16**) gives

$$\frac{p_+}{1-p_+} k_3 = \frac{-(k_3 + k_5)^2 + k_5(k_+ + k_-) + k_3(k_+ + k_-) - k_+ k_-}{k_3}$$

$$0 = {k_3}^2 \left(1 + \frac{p_+}{1-p_+}\right) + k_3(+2k_5 - k_+ - k_-) + k_+ k_- - k_5(k_+ + k_-) + {k_5}^2$$

And $1 + \frac{p_+}{1-p_+} = \frac{1}{1-p_+}$, thus

$$k_3 = \frac{1}{2}(k_+ + k_- - 2k_5)(1 - p_+) \pm \frac{1}{2}(1 - p_+) \sqrt{(k_+ + k_- - 2k_5)^2 - \frac{4(k_+ - k_5)(k_- - k_5)}{1 - p_+}}$$

And we already derived relations for $k_5$ (**Equation S15**) and $k_4$ (**Equation S17**) in terms of $p_+$, $k_+$, $k_-$,$k_3$, so this equation gives us a complete set of conversion relations.

$$k_5 = p_+ k_+ + (1 - p_+) k_-$$

$$k_3 = \frac{1}{2}(k_+ + k_- - 2k_5)(1 - p_+) + \frac{1}{2}(1 - p_+) \sqrt{(k_+ + k_- - 2k_5)^2 - \frac{4(k_+ - k_5)(k_- - k_5)}{1 - p_+}}$$

$$k_4 = k_5 - \frac{(k_+ - k_5)(k_5 - k_-)}{k_3}$$

To ensure $k_3, k_{-3}, k_4, k_5 \geq 0$ for this assumption, we need to take the expression for $k_3$ with the plus-sign before the second term and the expression under the square root should be positive or zero. For this second condition, we must have $k_+ \geq k_5 \geq k_-$ and $p_+ \geq 0$, which boils down to $p_+ \geq 0$ and $k_+ \geq k_-$ and these conditions are per definition true for a double exponential fit.

Additionally, to ensure $k_4 > 0$ we have the boundary

$k_4 = k_5 - \frac{(k_+ - k_5)(k_5 - k_-)}{k_3} > 0$ thus $k_3 > \frac{(k_+ - k_5)(k_5 - k_-)}{k_5}$

Filling in this boundary in the expression for $k_3$ and using the expression for $k_5$ (**Equation S15**) we can get the upper boundary on $p_+$

$$p_+ < \frac{k_+ k_+}{k_5(k_+ + k_- - k_5)} = \frac{\sqrt{k_+ k_+}}{k_+ - k_-}$$

By evaluating this boundary for the double exponential fit parameters obtained, we could directly argue whether this model assumption would mathematically work or not.

**Model 3, Assumption 2**: **No reverse reaction from $RP_O$ to $RP_I$**

For the assumption $k_{-3} = 0$, we find from the general expression for $k_{-3}$ (**Equation S16**) that $k_+ = k_3 + k_5$ or $k_- = k_3 + k_5$. Considering $k_3 + k_5 > k_4$ as we expect unbinding to be faster from $RP_I$ then from $RP_O$, we take $k_+ = k_3 + k_5$ and obtain

$$k_4 = k_5 - \frac{((k_3 + k_5) - k_5)(k_5 - k_-)}{k_3} = k_-$$

Taking into account the expression for $k_5$ (**Equation S15**), we have a complete set of conversion relations.

$k_5 = p_+ k_+ + (1 - p_+)k_-$

$k_4 = k_-$

$k_3 = k_+ - k_5$

For $k_3 \geq 0$ we need to have $k_+ \geq k_5$, which boils down to $p_+ \geq 0$ and $k_+ \geq k_-$ and these conditions are per definition true for a double exponential fit.

**Model 3, Assumption 3: dissociation from only $RP_I$**

Alternatively, we can assume there is no dissociation from $RP_O$, thus $k_4 = 0$. Under this assumption we get $k_5 = \frac{(k_+ - k_5)(k_5 - k_-)}{k_3}$ from **Equation S17** and $k_+ k_- = k_5 k_{-3}$ from **Equation S13**, thus we have

$k_5 = p_+ k_+ + (1 - p_+)k_-$

$k_3 = \frac{(k_+ - k_5)(k_5 - k_-)}{k_5}$

$k_{-3} = \frac{k_+ k_-}{k_5}$

In this case, we must have $k_+ \geq k_5 \geq k_-$ to ensure $k_3, k_{-3}, k_4, k_5 \geq 0$, which means $p_+ \geq 0$ and $k_+ \geq k_-$ and these conditions are per definition true for a double exponential fit. The evaluation of the kinetic rates calculated from the two exponential fits for the different assumptions to **Model 3** demonstrated that **Assumption 3** was the best proposed model for the OS disassembly times (**Results, Table 1**).

**Supplementary Notes 2: Kinetic description of single exponential OS dwell time distributions**

Using Model 3, Assumption 3 to describe the OS dwell times (**Table 1**), the OS dwell times distributions described by a single exponential probability distribution function report on the direct dissociation from $RP_I$ with a rate $k_5$, i.e. the holo did not transition to $RP_O$.

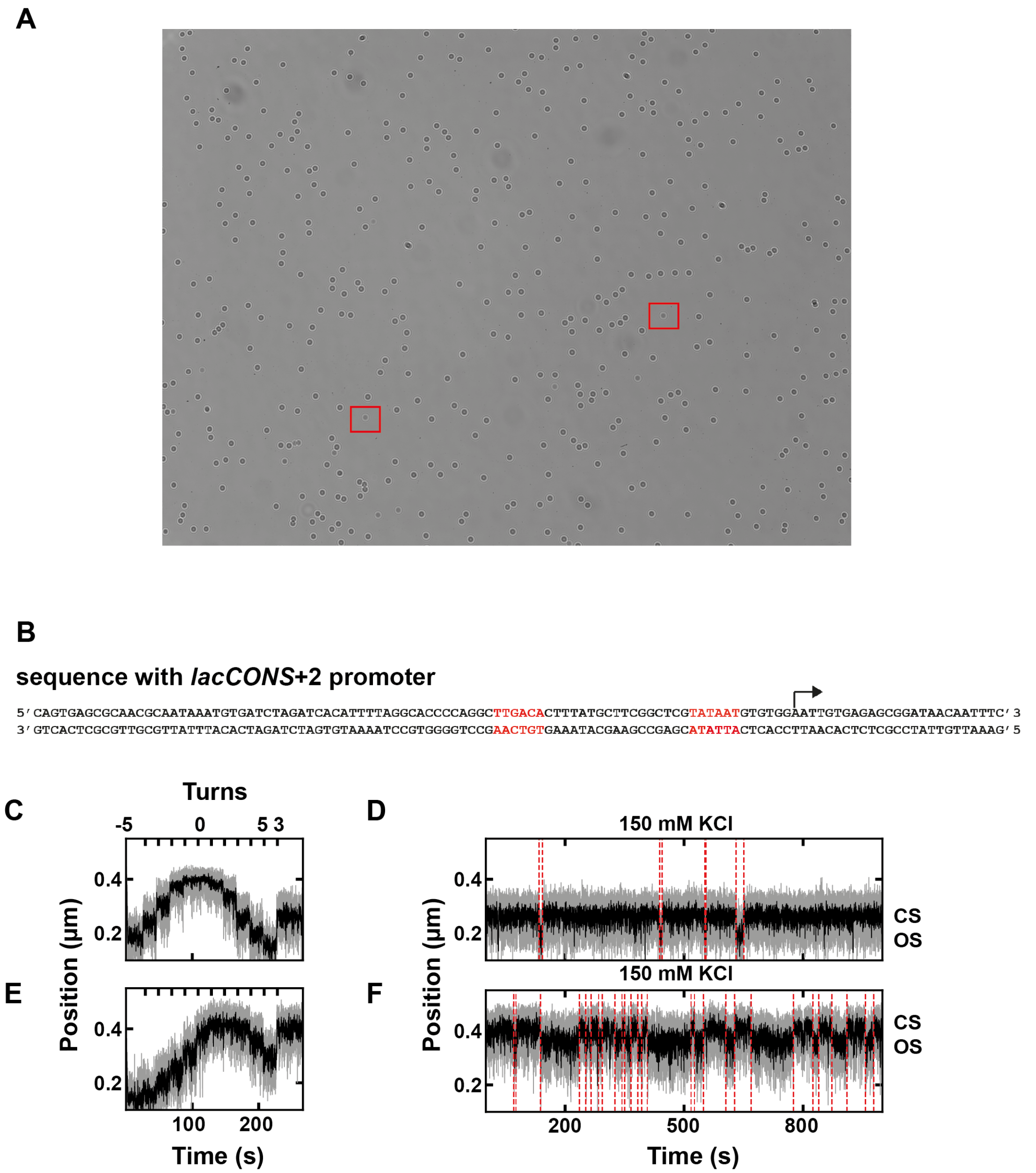


**Supplementary Figure 1. Field of view, DNA sequence and effects of torque on open complex dynamics. (A)** Typical field of view from a high throughput magnetic tweezers experiment, where 1 µm diameter MyOne magnetic beads (dark circles) are tethered to the glass surface by a ~1.4 kbp coilable DNA construct (**Material and Methods**). The polystyrene reference beads (1.1 µm diameter) are surrounded by a red square. **(B)** *lacCONS*+2 promoter sequence that is inserted in the dsDNA construct used in the MT experiment. The -10 element and -35 element are indicated in red, and the transcription start site and the holo transcription direction are indicated by an arrow. **(C)** The rotation extension of a tethered magnetic bead that is centered at zero turn and the measurement position (+3 turns) are beyond the plectonemic transition, i.e. in the constant torque regime. **(D)** Open complex dynamics for the tether calibrated in (C) with 150 mM KCl and 10 nM holo. **(E)** The rotation extension of a tethered magnetic bead performed in the same experiment as in (C), where the measurement position (+3 turns) precedes the plectonemic transition, i.e. at a lower torque than in (C). **(F)** Open complex dynamics for the tether calibrated in (E). The dashed red vertical lines indicate the detected transitions in (D) and (F) from the change-point analysis (**Materials and Methods**).

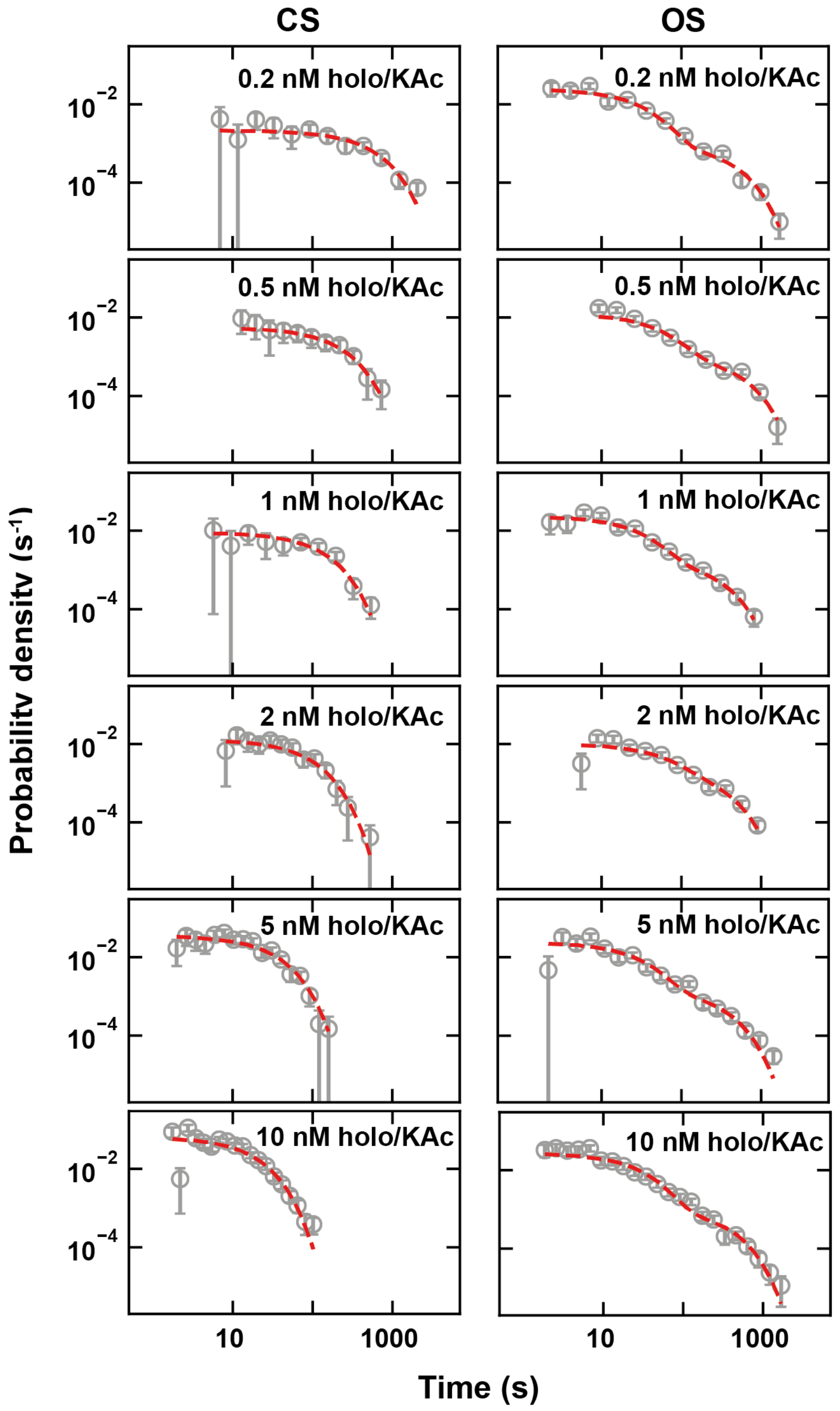


**Supplementary Figure 2. CS and OS dwell times distribution as a function of holo concentration.** Dwell time distributions of the open and closed states, i.e. OS and CS respectively, in 150 mM KAc at the holo concentration indicated in the panels. The dashed red line is either mono-exponential or bi-exponential MLE fit. Error bars are two standard deviations extracted from 1000 bootstraps.

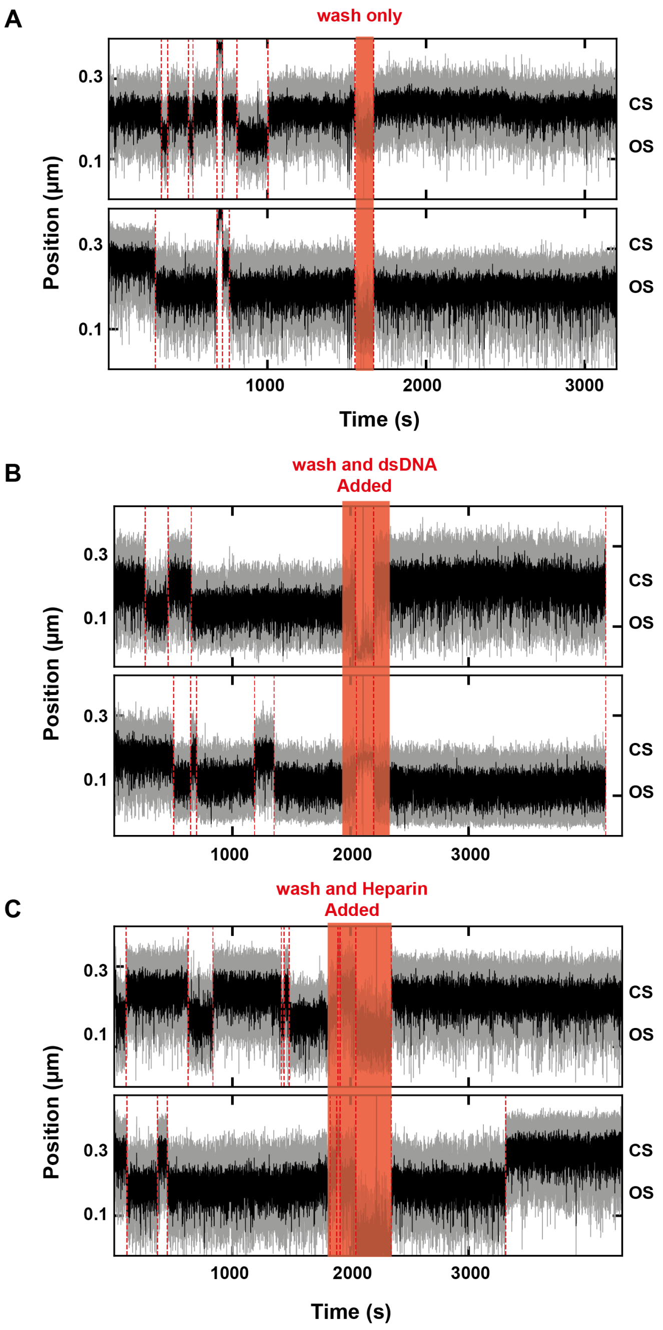


**Supplementary Figure 3. The transition from open to closed state releases free the bound holo from the DNA. (A)** Two representative traces showing the open complex dynamics for 0.5 nM holo and 150 mM of KGlu. The flow chamber is rinsed with 0.4 ml of reaction buffer at ~1800 s, and nothing is added afterwards. **(B)** Same experiment as in (A), and 0.2 ml of ~10 nM competing *lacCONS*+2 DNA promoter is then flushed after the flow chamber is rinsed with 0.2 ml of reaction buffer. **(C)** Same experiments as in (A), but 0.2 ml of 100 µg/ml heparin is added instead of competing DNA promoter after the flow chamber is rinsed with 0.2 ml of reaction buffer.

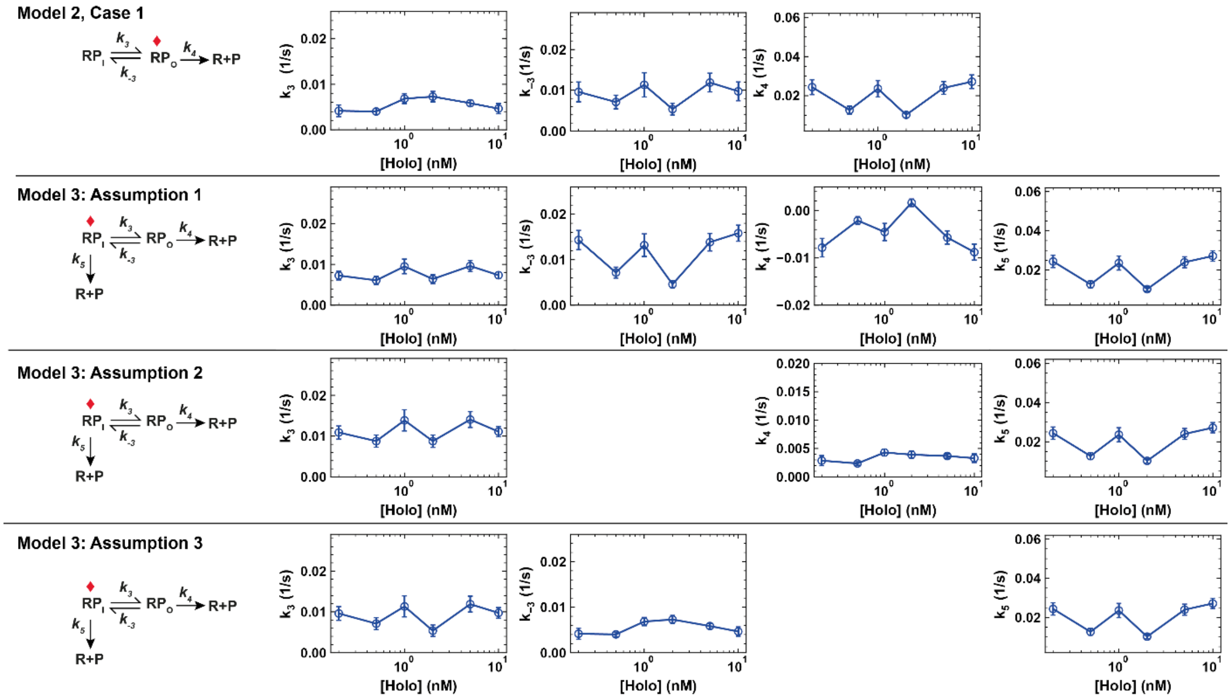


**Supplementary Figure 4. Comparison of microscopic rates determined using the different models described in Table 1.** Plots of the microscopic rates for Model 2 and Model 3. The error bars are one standard deviation extracted from error propagation as described in **Materials and Methods**.

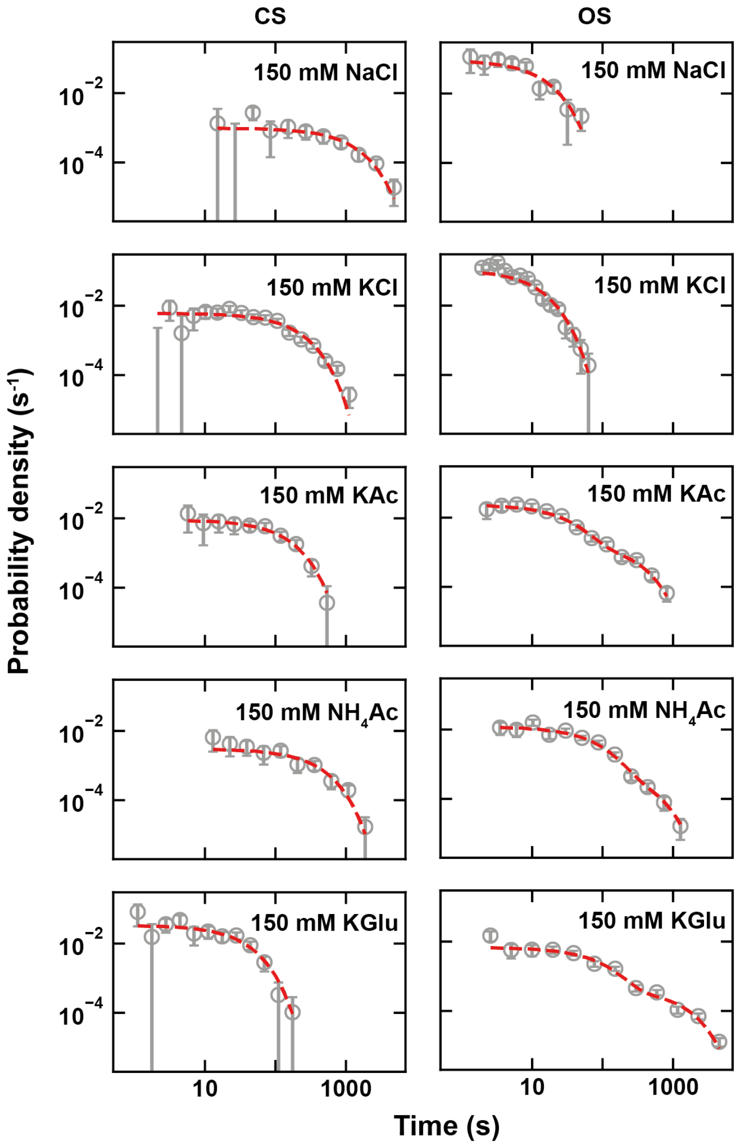


**Supplementary Figure 5. CS and OS dwell times distribution as a function of monovalent salt types.** Dwell time distributions of the open (OS) and closed (CS) states for different monovalent salts at the same concentration. 10 nM holo was used for KCl and NaCl, while 1 nM holo was used in the other conditions. The dashed red line is the either mono-exponential or bi-exponential MLE fit. Error bars are two standard deviations extracted from 1000 bootstraps.

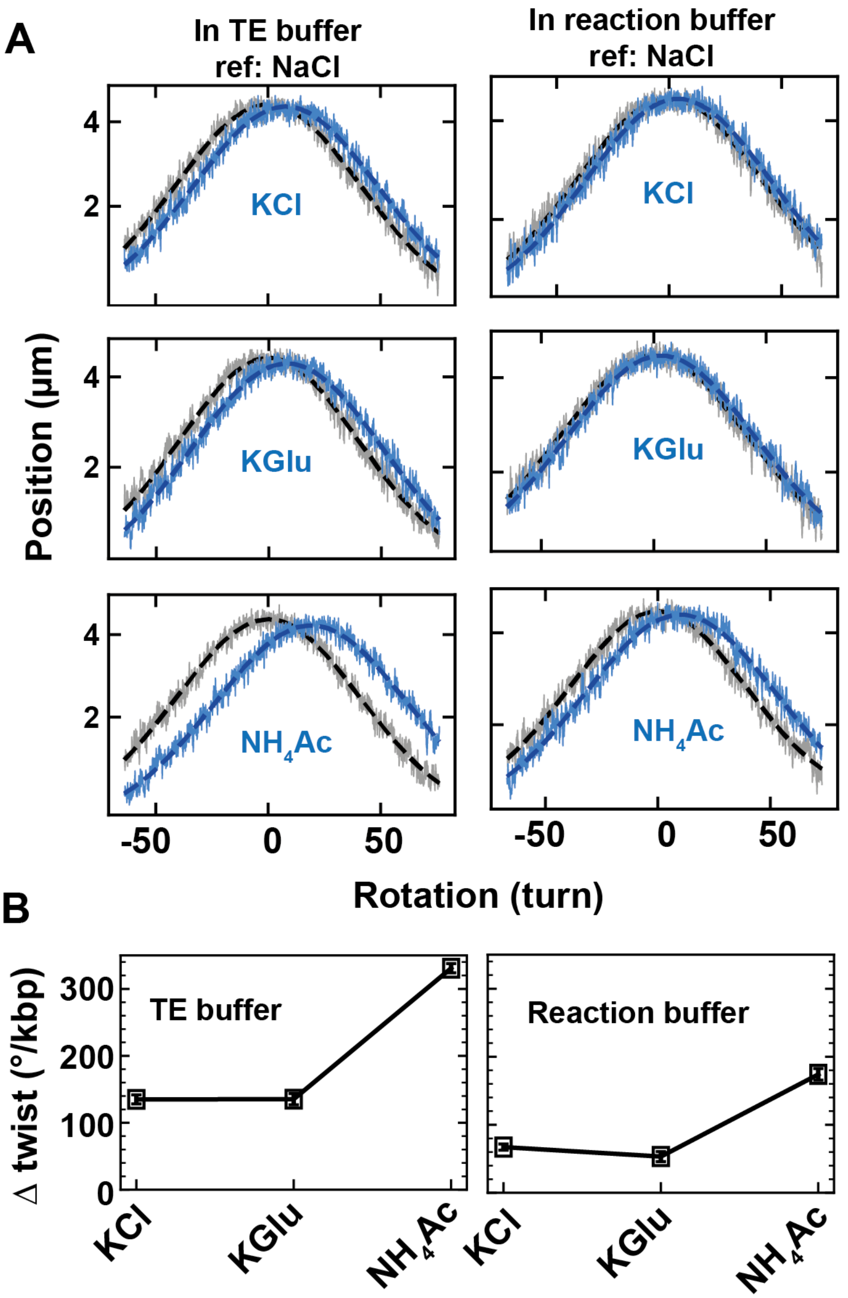


**Supplementary Figure 6. The cation nature affects the DNA helical twist. (A)** Extension as a function of DNA rotation (extension-rotation) for different monovalent salts at 150 mM concentration, either in TE buffer (left) or holo reaction buffer (right). The reference extension-rotation in NaCl is in black and the one measured in the indicated monovalent salt is in blue. The dashed lines are their respective Gaussian fit. **(B)** Change in DNA helical twist (Δtwist) per degree of rotation and per kilo base pair in comparison of the reference measurement performed with 150 mM NaCl in either TE buffer (left) or reaction buffer (right).

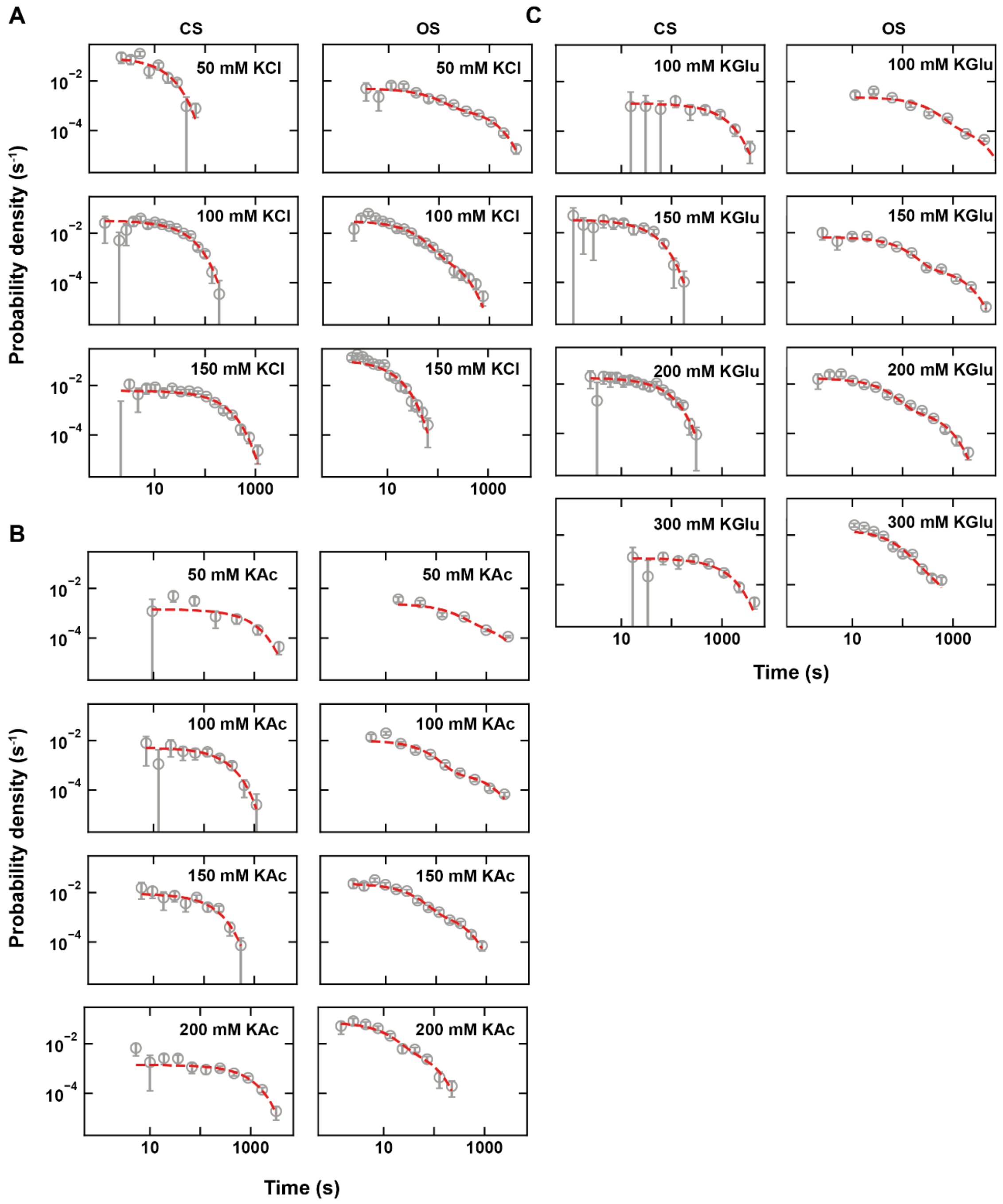


**Supplementary Figure 7. CS and OS dwell times distribution as a function of anions concentration.** Dwell time distributions of the open and closed states, i.e. OS and CS respectively, for **(A)** KCl, **(B)** KAc and **(C)** KGlu at the concentration indicated in the panels. 10 nM holo was used for KCl and 1 nM holo was used for other anions. The dashed red line is the either mono-exponential or bi-exponential MLE fit. Error bars are two standard deviations extracted from 1000 bootstraps.

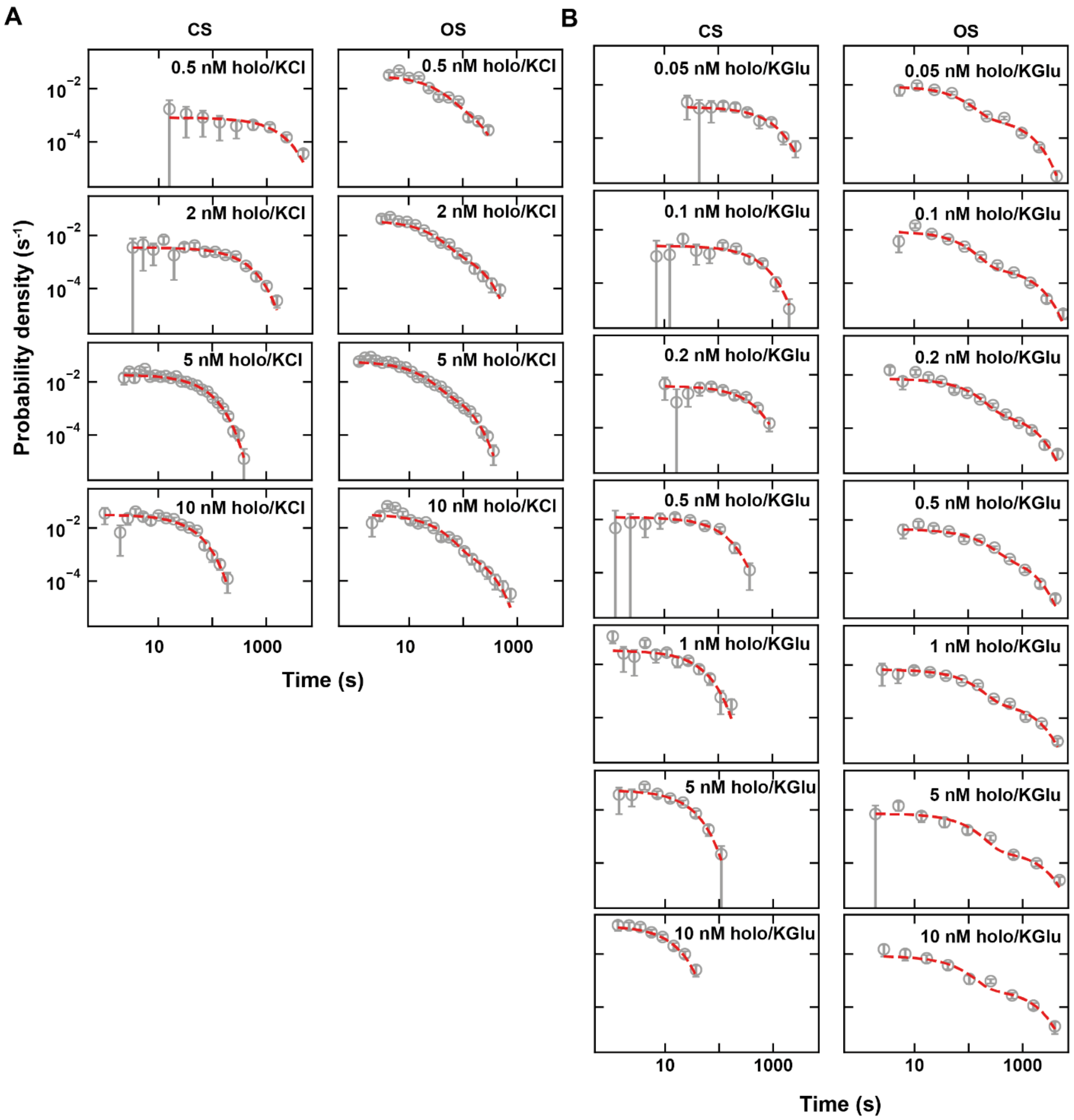


**Supplementary Figure 8. CS and OS dwell times distribution as a function of holo concentration.** Dwell time distributions of the open and closed states, i.e. OS and CS respectively, in (A) 100 mM KCl and (B) 150 mM KGlu at the holo concentration indicated in the panels. The dashed red line is either mono-exponential or bi-exponential MLE fit. Error bars are two standard deviations extracted from 1000 bootstraps.

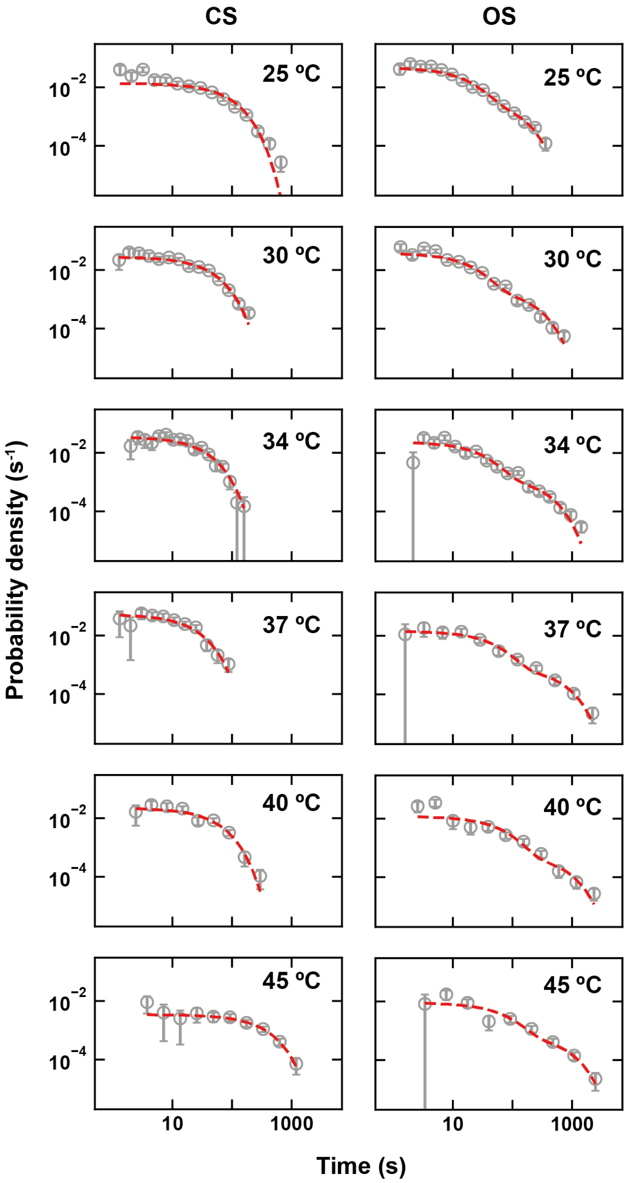


**Supplementary Figure 9. Effect of temperature on the open complex dynamics.** Dwell time distributions of the OS and CS at the indicated temperature, performed with in 150 mM KAc and 5 nM holo. The dashed red line is either a single exponential (CS) or a double exponential (OS) MLE fit. Error bars are two standard deviations extracted from 1000 bootstraps.

**Table S1:** Experimental conditions and their related parameters extracted from MLE procedures. The rate constants $k_3$, $k_{-3}$ and $k_5$ were extracted according to Model 3, Assumption 3.

| Experiment Conditions | | | T ('C) | Used in figures | Number of traces | | | | Close state dwell time | | | Open state dwell time | | | | | | |
|---|---|---|---|---|---|---|---|---|---|---|---|---|---|---|---|---|---|---|
| Salt | [Salt] (mM) | [Holo] (nM) | | | Total | Active | active frac. | 95% CI | Dwell times | k_open (1/s) | SD (1/s) | Dwell times | k3 (1/s) | SD (1/s) | k-3 (1/s) | SD (1/s) | k5 (1/s) | SD (1/s) |
| KCl | 50 | 10 | 25 | 3, S4 | 77 | 75 | 0.97 | 0.04 | 236 | 0.0894 | 0.0055 | 258 | 0.008 | 0.006 | 0.0036 | 0.0010 | 0.005 | 0.003 |
| | 100 | | | 3, 4, S4, S5 | 37 | 34 | 0.92 | 0.09 | 926 | 0.0320 | 0.0010 | 922 | 0.007 | 0.003 | 0.0090 | 0.0022 | 0.032 | 0.005 |
| | 150 | | | 2, 3, S1, S2, S4 | 39 | 38 | 0.97 | 0.05 | 1031 | 0.0061 | 0.0002 | 1030 | - | - | - | - | 0.109 | 0.003 |
| | 100 | 10 | | 3, 4, S4, S5 | 37 | 34 | 0.92 | 0.09 | 926 | 0.0320 | 0.0010 | 922 | 0.007 | 0.003 | 0.0090 | 0.0022 | 0.032 | 0.005 |
| | | 5 | | 4, S5 | 54 | 53 | 0.98 | 0.04 | 3639 | 0.0187 | 0.0003 | 3641 | 0.020 | 0.004 | 0.0244 | 0.0014 | 0.059 | 0.007 |
| | | 2 | | 4, S5 | 59 | 56 | 0.95 | 0.06 | 602 | 0.0035 | 0.0002 | 579 | 0.012 | 0.004 | 0.0129 | 0.0023 | 0.037 | 0.006 |
| | | 0.5 | | 4, S5 | 48 | 41 | 0.85 | 0.10 | 162 | 0.0008 | 0.0001 | 137 | 0.008 | 0.004 | 0.0146 | 0.0029 | 0.031 | 0.007 |
| KAc | 50 | 1 | 25 | 3, S4 | 60 | 57 | 0.95 | 0.06 | 101 | 0.0014 | 0.0002 | 78 | 0.002 | 0.001 | 0.0015 | 0.0003 | 0.002 | 0.001 |
| | 100 | | | 3, S4 | 36 | 36 | 1.00 | 0.00 | 133 | 0.0052 | 0.0005 | 125 | 0.008 | 0.005 | 0.0021 | 0.0019 | 0.010 | 0.005 |
| | 150 | | | 2, 3, S2, S4 | 29 | 27 | 0.93 | 0.09 | 207 | 0.0090 | 0.0006 | 196 | 0.011 | 0.003 | 0.0068 | 0.0011 | 0.024 | 0.004 |
| | 200 | | | 3, S4 | 61 | 59 | 0.97 | 0.04 | 453 | 0.0014 | 0.0001 | 418 | 0.030 | 0.006 | 0.0272 | 0.0031 | 0.076 | 0.009 |
| | 150 | 10 | | 1, 4, S5 | 57 | 56 | 0.98 | 0.04 | 1479 | 0.0649 | 0.0017 | 1455 | 0.010 | 0.002 | 0.0047 | 0.0011 | 0.027 | 0.003 |
| | | 5 | | 4, 5, 6, S5, S7 | 51 | 49 | 0.96 | 0.05 | 770 | 0.0353 | 0.0012 | 781 | 0.012 | 0.002 | 0.0059 | 0.0007 | 0.024 | 0.003 |
| | | 2 | | 4, S5 | 46 | 44 | 0.96 | 0.06 | 286 | 0.0129 | 0.0007 | 306 | 0.005 | 0.001 | 0.0073 | 0.0011 | 0.010 | 0.001 |
| | | 1 | | 2, 3, S2, S4 | 29 | 27 | 0.93 | 0.09 | 207 | 0.0090 | 0.0006 | 196 | 0.011 | 0.003 | 0.0068 | 0.0011 | 0.024 | 0.004 |
| | | 0.5 | | 4, S5 | 38 | 37 | 0.97 | 0.05 | 164 | 0.0056 | 0.0004 | 176 | 0.007 | 0.002 | 0.0040 | 0.0006 | 0.013 | 0.002 |
| | | 0.2 | | 4, S5 | 46 | 43 | 0.94 | 0.07 | 273 | 0.0026 | 0.0001 | 260 | 0.010 | 0.002 | 0.0042 | 0.0013 | 0.024 | 0.004 |
| KGlu | 100 | 1 | 25 | 3, S4 | 50 | 48 | 0.96 | 0.05 | 102 | 0.0013 | 0.0001 | 104 | 0.001 | 0.001 | 0.0006 | 0.0002 | 0.002 | 0.001 |
| | 150 | | | 2, 3, S2, S4 | 47 | 45 | 0.96 | 0.06 | 243 | 0.0338 | 0.0023 | 230 | 0.005 | 0.002 | 0.0016 | 0.0019 | 0.007 | 0.002 |
| | 200 | | | 3, S4 | 49 | 49 | 1.00 | 0.00 | 481 | 0.0179 | 0.0008 | 466 | 0.010 | 0.003 | 0.0039 | 0.0008 | 0.017 | 0.005 |
| | 300 | | | 3, S4 | 43 | 40 | 0.93 | 0.08 | 206 | 0.0012 | 0.0001 | 193 | 0.003 | 0.002 | 0.0055 | 0.0040 | 0.017 | 0.005 |
| | 150 | 0.05 | | 4, S5 | 53 | 52 | 0.98 | 0.04 | 168 | 0.0015 | 0.0001 | 158 | 0.008 | 0.003 | 0.0023 | 0.0016 | 0.009 | 0.003 |
| | | 0.1 | | 4, S5 | 65 | 64 | 0.99 | 0.03 | 251 | 0.0025 | 0.0002 | 245 | 0.007 | 0.003 | 0.0016 | 0.0038 | 0.009 | 0.003 |
| | | 0.2 | | 4, S5 | 49 | 47 | 0.96 | 0.06 | 268 | 0.0038 | 0.0002 | 267 | 0.004 | 0.003 | 0.0016 | 0.0012 | 0.007 | 0.004 |
| | | 0.5 | | 2, 4S, S6 | 64 | 61 | 0.95 | 0.06 | 272 | 0.0124 | 0.0008 | 263 | 0.002 | 0.002 | 0.0018 | 0.0007 | 0.005 | 0.002 |
| | | 1 | | 2, 3, S2, S4 | 47 | 45 | 0.96 | 0.06 | 243 | 0.0338 | 0.0023 | 230 | 0.005 | 0.002 | 0.0016 | 0.0019 | 0.007 | 0.002 |
| | | 5 | | 4, S5 | 56 | 54 | 0.96 | 0.05 | 322 | 0.0554 | 0.0035 | 335 | 0.005 | 0.001 | 0.0012 | 0.0002 | 0.007 | 0.002 |
| | | 10 | | 4, S5 | 58 | 56 | 0.97 | 0.04 | 318 | 0.1161 | 0.0058 | 335 | 0.009 | 0.002 | 0.0020 | 0.0003 | 0.008 | 0.002 |
| NaCl | 150 | 10 | 25 | 2, S2 | 53 | 13 | 0.25 | 0.12 | 66 | 0.0010 | 0.0001 | 54 | - | - | - | - | 0.092 | 0.008 |
| NH4Ac | 150 | 1 | 25 | 2, S2 | 45 | 43 | 0.96 | 0.06 | 179 | 0.0030 | 0.0002 | 169 | 0.002 | 0.002 | 0.0036 | 0.0031 | 0.012 | 0.004 |
| KAc | 150 | 5 | 25 | 4, 5, 6, S5, S7 | 41 | 40 | 0.98 | 0.04 | 807 | 0.0137 | 0.0005 | 1448 | 0.016 | 0.005 | 0.0153 | 0.0012 | 0.047 | 0.008 |
| | | | 30 | 5, 6, S7 | 44 | 47 | 0.94 | 0.07 | 1587 | 0.0280 | 0.0008 | 1647 | 0.017 | 0.003 | 0.0090 | 0.0016 | 0.038 | 0.004 |
| | | | 34 | 5, 6, S7 | 51 | 49 | 0.96 | 0.05 | 770 | 0.0350 | 0.0012 | 781 | 0.012 | 0.002 | 0.0059 | 0.0007 | 0.024 | 0.003 |
| | | | 37 | 5, 6, S7 | 54 | 53 | 0.98 | 0.04 | 393 | 0.0520 | 0.0025 | 409 | 0.008 | 0.002 | 0.0033 | 0.0004 | 0.015 | 0.003 |
| | | | 40 | 5, 6, S7 | 46 | 45 | 0.98 | 0.04 | 383 | 0.0220 | 0.0013 | 380 | 0.005 | 0.002 | 0.0024 | 0.0005 | 0.012 | 0.003 |
| | | | 45 | 5, 6, S7 | 49 | 49 | 1.00 | 0.00 | 284 | 0.0040 | 0.0002 | 278 | 0.005 | 0.002 | 0.0027 | 0.0004 | 0.009 | 0.002 |

**Table S2:** Experimental conditions and their parameters directly obtained from MLE fits

| Experiment Conditions | | | T (°C) | Used in figures | Number of traces | | | | Close state dwell time | | | Open state dwell time | | | | | | |
|---|---|---|---|---|---|---|---|---|---|---|---|---|---|---|---|---|---|---|
| Salt | [Salt] (mM) | [Holo] (nM) | | | Total | Active | Active frac. | 95% CI | Dwell times | k_open (1/s) | SD (1/s) | Dwell times | k_(1/s) | SD (1/s) | k+ (1/s) | SD (1/s) | p+ (1/s) | SD (1/s) |
| KCl | 50 | 10 | 25 | 3, S4 | 77 | 75 | 0.97 | 0.04 | 236 | 0.0894 | 0.0055 | 258 | 0.016 | 0.008 | 0.0012 | 0.0002 | 0.73 | 0.08 |
| | 100 | | | 3, 4, S4, S5 | 37 | 34 | 0.92 | 0.09 | 926 | 0.0320 | 0.0010 | 922 | 0.041 | 0.006 | 0.0070 | 0.0016 | 0.27 | 0.08 |
| | 150 | | | 2, 3, S1, S2, S4 | 39 | 38 | 0.97 | 0.05 | 1031 | 0.0061 | 0.0002 | 1030 | 0.109 | 0.003 | - | - | - | - |
| | 100 | 10 | | 3, 4, S4, S5 | 37 | 34 | 0.92 | 0.09 | 926 | 0.0320 | 0.0010 | 922 | 0.041 | 0.006 | 0.0070 | 0.0016 | 0.27 | 0.08 |
| | | 5 | | 4, S5 | 54 | 53 | 0.98 | 0.04 | 3639 | 0.0187 | 0.0003 | 3641 | 0.086 | 0.012 | 0.0166 | 0.0008 | 0.40 | 0.03 |
| | | 2 | | 4, S5 | 59 | 56 | 0.95 | 0.06 | 602 | 0.0035 | 0.0002 | 579 | 0.053 | 0.008 | 0.0089 | 0.0015 | 0.37 | 0.07 |
| | | 0.5 | | 4, S5 | 48 | 41 | 0.85 | 0.10 | 162 | 0.0008 | 0.0001 | 137 | 0.044 | 0.009 | 0.0104 | 0.0019 | 0.38 | 0.11 |
| KAc | 50 | 1 | 25 | 3, S4 | 60 | 57 | 0.95 | 0.06 | 101 | 0.0014 | 0.0002 | 78 | 0.005 | 0.001 | 0.0007 | 0.0001 | 0.63 | 0.08 |
| | 100 | | | 3, S4 | 36 | 36 | 1.00 | 0.00 | 133 | 0.0052 | 0.0005 | 125 | 0.019 | 0.008 | 0.0011 | 0.0011 | 0.49 | 0.09 |
| | 150 | | | 2, 3, S2, S4 | 29 | 27 | 0.93 | 0.09 | 207 | 0.0090 | 0.0006 | 196 | 0.037 | 0.006 | 0.0043 | 0.0006 | 0.42 | 0.06 |
| | 200 | | | 3, S4 | 61 | 59 | 0.97 | 0.04 | 453 | 0.0014 | 0.0001 | 418 | 0.116 | 0.013 | 0.0179 | 0.0018 | 0.40 | 0.05 |
| | 150 | 10 | | 1, 4, S5 | 57 | 56 | 0.98 | 0.04 | 1479 | 0.0649 | 0.0017 | 1455 | 0.038 | 0.004 | 0.0033 | 0.0008 | 0.32 | 0.07 |
| | | 5 | | 4, 5, 6, S5, S7 | 51 | 49 | 0.96 | 0.05 | 770 | 0.0353 | 0.0012 | 781 | 0.038 | 0.005 | 0.0037 | 0.0004 | 0.41 | 0.05 |
| | | 2 | | 4, S5 | 46 | 44 | 0.96 | 0.06 | 286 | 0.0129 | 0.0007 | 306 | 0.019 | 0.003 | 0.0039 | 0.0006 | 0.58 | 0.06 |
| | | 1 | | 2, 3, S2, S4 | 29 | 27 | 0.93 | 0.09 | 207 | 0.0090 | 0.0006 | 196 | 0.037 | 0.006 | 0.0043 | 0.0006 | 0.42 | 0.06 |
| | | 0.5 | | 4, S5 | 38 | 37 | 0.97 | 0.05 | 164 | 0.0056 | 0.0004 | 176 | 0.021 | 0.003 | 0.0024 | 0.0003 | 0.46 | 0.06 |
| | | 0.2 | | 4, S5 | 46 | 43 | 0.94 | 0.07 | 273 | 0.0026 | 0.0001 | 260 | 0.035 | 0.005 | 0.0029 | 0.0009 | 0.34 | 0.07 |
| KGlu | 100 | 1 | 25 | 3, S4 | 50 | 48 | 0.96 | 0.05 | 102 | 0.0013 | 0.0001 | 104 | 0.003 | 0.002 | 0.0004 | 0.0001 | 0.46 | 0.09 |
| | 150 | | | 2, 3, S2, S4 | 47 | 45 | 0.96 | 0.06 | 243 | 0.0338 | 0.0023 | 230 | 0.013 | 0.004 | 0.0009 | 0.0011 | 0.49 | 0.07 |
| | 200 | | | 3, S4 | 49 | 49 | 1.00 | 0.00 | 481 | 0.0179 | 0.0008 | 466 | 0.029 | 0.007 | 0.0023 | 0.0005 | 0.44 | 0.07 |
| | 300 | | | 3, S4 | 43 | 40 | 0.93 | 0.08 | 206 | 0.0012 | 0.0001 | 193 | 0.020 | 0.005 | 0.0045 | 0.0034 | 0.24 | 0.15 |
| | 150 | 0.05 | | 4, S5 | 53 | 52 | 0.98 | 0.04 | 168 | 0.0015 | 0.0001 | 158 | 0.018 | 0.005 | 0.0012 | 0.0008 | 0.53 | 0.09 |
| | | 0.1 | | 4, S5 | 65 | 64 | 0.99 | 0.03 | 251 | 0.0025 | 0.0002 | 245 | 0.016 | 0.004 | 0.0009 | 0.0022 | 0.48 | 0.06 |
| | | 0.2 | | 4, S5 | 49 | 47 | 0.96 | 0.06 | 268 | 0.0038 | 0.0002 | 267 | 0.012 | 0.006 | 0.0010 | 0.0008 | 0.42 | 0.08 |
| | | 0.5 | | 2, 4S, S6 | 64 | 61 | 0.95 | 0.06 | 272 | 0.0124 | 0.0008 | 263 | 0.008 | 0.003 | 0.0011 | 0.0004 | 0.47 | 0.10 |
| | | 1 | | 2, 3, S2, S4 | 47 | 45 | 0.96 | 0.06 | 243 | 0.0338 | 0.0023 | 230 | 0.013 | 0.004 | 0.0009 | 0.0011 | 0.49 | 0.07 |
| | | 5 | | 4, S5 | 56 | 54 | 0.96 | 0.05 | 322 | 0.0554 | 0.0035 | 335 | 0.013 | 0.003 | 0.0007 | 0.0001 | 0.46 | 0.04 |
| | | 10 | | 4, S5 | 58 | 56 | 0.97 | 0.04 | 318 | 0.1161 | 0.0058 | 335 | 0.018 | 0.004 | 0.0009 | 0.0001 | 0.57 | 0.04 |
| NaCl | 150 | 10 | 25 | 2, S2 | 53 | 13 | 0.25 | 0.12 | 66 | 0.0010 | 0.0001 | 54 | 0.092 | 0.008 | - | - | - | - |
| NH4Ac | 150 | 1 | 25 | 2, S2 | 45 | 43 | 0.96 | 0.06 | 179 | 0.0030 | 0.0002 | 169 | 0.015 | 0.004 | 0.0029 | 0.0026 | 0.25 | 0.14 |
| KAc | 150 | 5 | 25 | 4, 5, 6, S5, S7 | 41 | 40 | 0.98 | 0.04 | 807 | 0.0137 | 0.0005 | 1448 | 0.068 | 0.013 | 0.0106 | 0.0007 | 0.36 | 0.04 |
| | | | 30 | 5, 6, S7 | 44 | 47 | 0.94 | 0.07 | 1587 | 0.0280 | 0.0008 | 1647 | 0.058 | 0.006 | 0.0059 | 0.0011 | 0.38 | 0.04 |
| | | | 34 | 5, 6, S7 | 51 | 49 | 0.96 | 0.05 | 770 | 0.0350 | 0.0012 | 781 | 0.038 | 0.005 | 0.0037 | 0.0004 | 0.41 | 0.05 |
| | | | 37 | 5, 6, S7 | 54 | 53 | 0.98 | 0.04 | 393 | 0.0520 | 0.0025 | 409 | 0.024 | 0.004 | 0.0020 | 0.0002 | 0.43 | 0.05 |
| | | | 40 | 5, 6, S7 | 46 | 45 | 0.98 | 0.04 | 383 | 0.0220 | 0.0013 | 380 | 0.018 | 0.005 | 0.0017 | 0.0003 | 0.35 | 0.07 |
| | | | 45 | 5, 6, S7 | 49 | 49 | 1.00 | 0.00 | 284 | 0.0040 | 0.0002 | 278 | 0.016 | 0.004 | 0.0016 | 0.0002 | 0.46 | 0.05 |

**Table S3:** Equilibrium constants determined from holo concentration dependent experiment fitting the data with **Equation 1**.

| Salt | $K_1$ ($\mu M^{-1}$) | $k_2$ ($s^{-1}$) |
|---|---|---|
| 100 mM KCl | 0.95 ± 4.96 | 3.0 ± 15.7 |
| 150 mM KAc | 60.1 ± 4.8 | 0.17 ± 0.01 |
| 150 mM KGlu | 6.5 ± 6.4 | 2.4 ± 2.3 |


1. Floyd, D.L., Harrison, S.C. and van Oijen, A.M. (2010) Analysis of kinetic intermediates in single-particle dwell-time distributions. *Biophysical journal*, **99**, 360-366.
2. Xie, S.N. (2001) Single-molecule approach to enzymology. *Single Mol*, **2**, 229-236.
